# A multidisciplinary approach towards identification of novel antibiotic scaffolds for *Acinetobacter baumannii*

**DOI:** 10.1101/306035

**Authors:** Satya Prathyusha Bhamidimarri, Michael Zahn, Jigneshkumar Dahyabhai Prajapati, Christian Schleberger, Sandra Söderholm, Jennifer Hoover, Josh West, Ulrich Kleinekathöfer, Dirk Bumann, Mathias Winterhalter, Bert van den Berg

## Abstract

Research efforts to discover potential new antibiotics for Gram-negative bacteria suffer from high attrition rates due to the synergistic action of efflux systems and the limited permeability of the outer membrane (OM). One potential strategy to overcome the OM permeability barrier is to identify small molecules that are natural substrates for abundant OM channels, and to use such compounds as scaffolds for the design of efficiently-permeating antibacterials. Here we present a multidisciplinary approach to identify such potential small-molecule scaffolds. Focusing on the pathogenic bacterium *Acinetobacter baumannii*, we use OM proteomics to identify DcaP as the most abundant channel under various conditions that are relevant for infection. High-resolution X-ray structure determination of DcaP surprisingly reveals a trimeric, porin-like structure and suggests that dicarboxylic acids are potential transport substrates. Electrophysiological experiments and allatom molecular dynamics simulations confirm this notion and provide atomistic information on likely permeation pathways and energy barriers for several small molecules, including a clinically-relevant β-lactamase inhibitor. Our study provides a general blueprint for the identification of molecular scaffolds that will inform the rational design of future antibacterials.

## Introduction

The outer membrane (OM) of Gram-negative bacteria forms a formidable barrier for both hydrophilic and hydrophobic molecules. For the uptake of nutrients and other low molecular weight hydrophilic compounds, water-filled channels in the OM act as entry pathways into the periplasm for small water-soluble molecules. In contrast to Enterobacteria that have relatively large and permanently open channels termed general porins that are not substrate specific, *Pseudomonas* and *Acinetobacter spp*. instead have substrate-specific channels with highly flexible loops, reducing the apparent pore size or even resulting in closed, gated pores. This provides an explanation for the very low permeability of the OM of these organisms (1–3) and makes them intrinsically resistant to many antibiotics. For such bacteria, it becomes even more important to understand on an atomic level how small molecules utilize the available channels for cellular entry. Such studies first require the identification of OM channels that are highly expressed *in vivo* during infection. A logical next step would be to identify the natural substrates taken up by such channels, and use these structures as scaffolds for the design of novel antibiotics, in combination with structural and computational studies to obtain atomistic understanding of permeation. Together, such an approach may lead to major advances, as it is now becoming clear that getting drugs into Gram-negative bacteria is a major challenge (4–7). One of the most problematic organisms is *Acinetobacter baumannii*, which has attracted attention due to its multi-drug resistance (MDR) (8). Currently available antibiotics effective against *Acinetobacter* include ampicillin, ticarcillin, meropenem and sulbactam (9, 10). Sulbactam is a β-lactamase inhibitor that has intrinsic antibacterial activity against *A. baumannii* by inactivating penicillin binding proteins (11, 12). However, until very recently, little was known about *A. baumannii* OM channels and their role in antibiotic uptake. This situation has changed considerably, and several OM channels of *A. baumannii* have now been characterised at least to some extent, including CarO, OccAB1 (OprD), OccAB2 (HcaE), OccAB3 (VanP), OccAB4 (BenP), OccAB5 and OmpA (1, 13, 14). Crystal structures are available for OccAB1-OccAB4 and CarO (13, 15).

In addition to OmpA (Omp38), CarO and OccAB1 (16, 17), another abundant OM protein from various antibiotic-resistant *A. baumannii* strains and biofilms is DcaP, first described by Parke et al. (18) in *Acinetobacter sp*. strain ADP1, and located in the *dca* (dicarboxylic acid) operon. Several other genes present in this region include *dcaA* coding for acyl-coA dehydrogenase, *dcaK* encoding an integral membrane protein belonging to the major facilitator superfamily, *dcaI* and *dcaJ* involved in the encoding of coA transferase subunit A and subunit B, respectively, and *dcaP* as a putative uptake channel. Relatively little is known about DcaP-like proteins, which appear to be confined among the *Moraxellaceae* family, and their roles in dicarboxylic acid uptake remain to be established.

Here, we use quantitative proteomics to show that DcaP is a highly abundant outer membrane protein in pathogenic *A. baumannii* strains during infection. The X-ray crystal structure shows that surprisingly, DcaP is a trimeric, “porin-like” OM protein. Additionally, DcaP has an N-terminal periplasmic domain that we predict via modelling to form a coiled-coil motif that might be involved in stabilizing the trimeric barrel. The abundance of the positively charged residues in the constriction region suggests a strong preference for negatively charged substrates such as succinates and phthalates. This notion is confirmed by standard electrophysiology experiments and reversal potential measurements, with the latter also providing translocation fluxes. The experiments are complemented with applied-field molecular dynamics (MD) and metadynamics simulations which confirm the permeation of the substrates through the DcaP channel. Finally, we show that DcaP is likely to be involved in the uptake of clinically-relevant negatively charged β-lactamase inhibitors, *e.g*. sulbactam and tazobactam.

## Results

A BLAST search suggests that DcaP is conserved among *Acinetobacter spp*. as well as closely related species (Fig. S1). DcaP was first identified in *Acinetobacter sp. strain ADP1* (also known as *Acinetobacter baylyi / strain ATCC 33305 / BD413 / ADP1)*, and was found to be located in a set of genes implicated to have a role in the uptake and metabolic pathway of dicarboxylic acids.(18) The focus of our structural studies, DcaP from *A. baumannii* AB307-0294 (ABBFA_000716), shares almost 30 % sequence identity with DcaP from *Acinetobacter sp. strain ADP1*. ABBFA_000716 is not located in an operon, but near a gene coding for lipid A ethanolamine phosphotransferase (ABBFA_000717, involved in polymyxin resistance), a gene encoding a protein belonging to major facilitator superfamily protein (ABBFA_000715) and ABBFA_000718 and ABBFA_000719, encoding transcriptional regulatory proteins qseA and qseB respectively (http://www.kegg.jp/dbget-bin/www_bget?abb:ABBFA_000716, (Fig. S2). Thus, the genetic context of ABBFA_000716 gives little clues about its function.

### Proteomic profiling identifies DcaP as an abundant OM protein during infection

To determine the abundance of DcaP (UniProt entry N9LF65) in *A. baumannii* ATCC 19606 in infected mouse and rat lung tissue, we employed a sensitive targeted proteomics approach on a high-resolution and accurate mass spectrometer with absolute quantification using heavy isotope-labelled reference peptides (Methods). As expected from the known, low-permeability OM of *A. baumannii*, there are no diffusion channels expressed at levels approaching those of OmpF and OmpC of *E. coli* (~10^5^ combined copies per cell). The results showed that Omp38 and an additional OmpA-like protein (A1S_0884) are the most abundant OMPs in *A. baumannii*, with 20,000 - 60,000 copies for OmpA. In both host species, DcaP was the third-most abundant OM protein, with 5,000 −15,000 DcaP copies per *A. baumannii* cell in mouse lung, and 2000-4000 copies per cell in rat lung (Fig. 1). Importantly, this indicates that DcaP is the most abundant OM diffusion channel *in vivo* because OmpA-like proteins mostly serve structural roles (19, 20); the putative large-pore conformer that may mediate non-specific diffusion of antibiotics (21) is present at only a few percent of the total (14), making it effectively a relatively minor species. In addition, no structural information is known for any large-pore OmpA conformer, precluding structure-based design of permeating compounds. In addition to DcaP, there are several other OMPs with uniform expression levels that seem high enough (> ~1000 copies/cell) for targeting by antibacterials.

**Figure 1.**
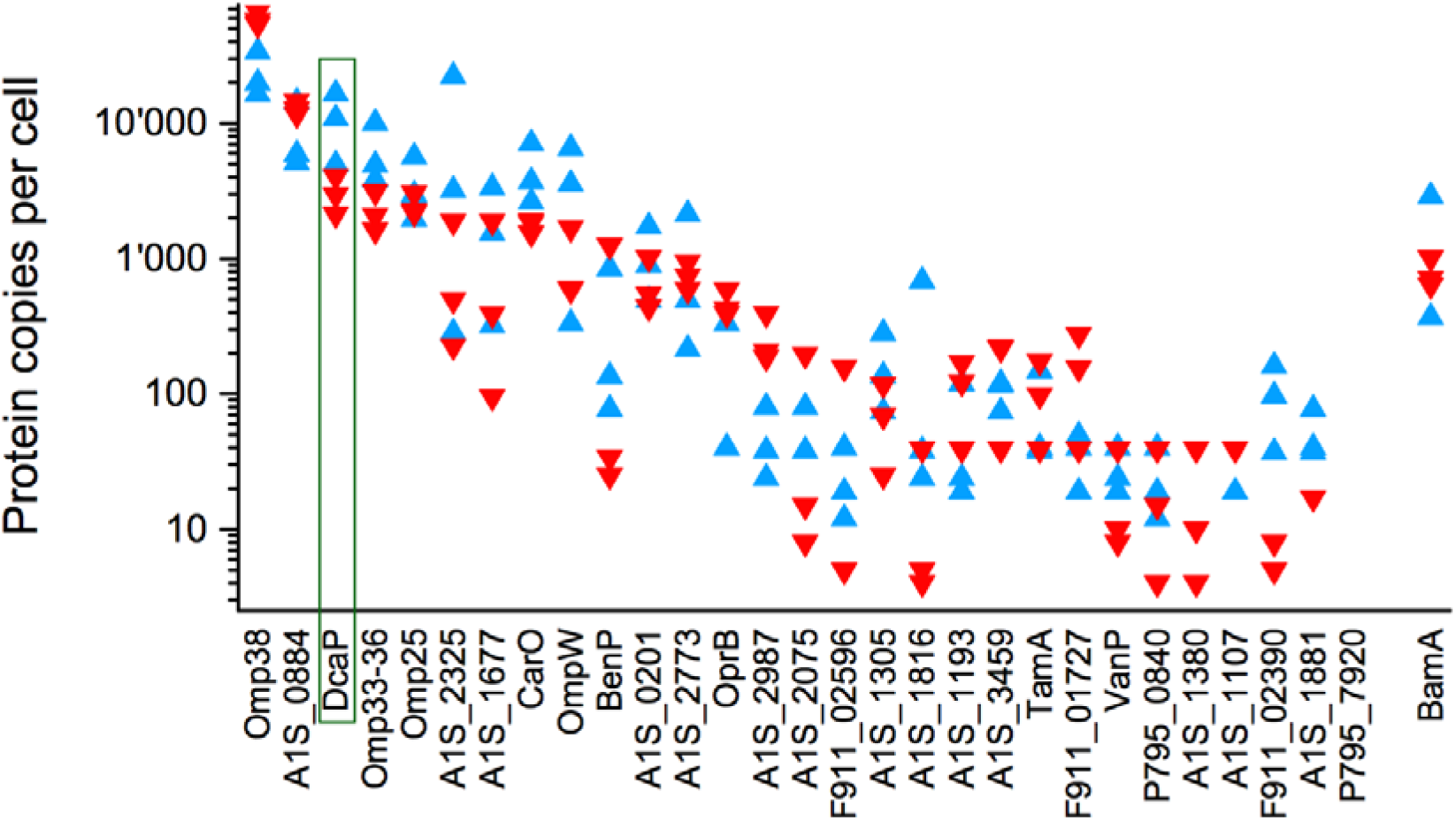
DcaP is highly abundant in *A. baumannii*. Copy numbers per cell for the known and putative OM diffusion channels determined by quantitative proteomics of infected mouse (blue symbols; n = 3) and rat (red symbols; n = 3) lung tissues. OmpA-like proteins, TamA and BamA are included as non-diffusion channels for comparison.

The first of these is Omp33-36, alternatively named Omp34, which is present at slightly lower levels compared to DcaP. No structure or detailed characterisation of this OMP is available, but it was recently predicted to form a 14-stranded barrel (22). Interestingly, Omp33-36 appears to be important for virulence of *A. baumannii* (23,24). It forms a water channel in *Xenopus* oocytes, but its proposed function as an OM aquaporin (23) seems highly unlikely given the presence of many other diffusion channels. Nevertheless, our proteomics data suggest that Omp33-36 is another interesting candidate for small-molecule targeting. Next, Omp25 and CarO, both of which are small proteins (~25 kDa), are two other OMPs present at high levels in all infected animals. A crystal structure is available only for CarO (15), and shows a very narrow, 8-stranded barrel that only allows uptake of very small molecules such as glycine (15). This is consistent with planar lipid bilayer experiments that show small ion conductance values (25). In the same study, Omp25 did not form channels (25). Thus, both CarO and Omp25 do not seem suitable for targeting by antibiotics, and the same most likely applies to OmpW, another 8-stranded β-barrel with a very narrow pore (26). Two other OMPs, A1S_2325 and A1S_1677, are expressed at high levels in some animals but much lower in others (Fig. 1), suggesting they are not very important for the bacterium. Both proteins have sizes (~35 kDa) compatible with diffusion channels but predictions (27) of 8- and 16-stranded barrels respectively suggest they are very different proteins. However, given the variation in expression levels during infection, we suggest that both these proteins are less promising candidates. All other potential diffusion channels of *A. baumannii* are expressed at low levels during infection. This includes OccAB1-4, which were recently characterised as 18-stranded barrels with relatively large pores, particularly OccAB1/OprD (13). Our proteomics data highlight the importance of assessing OMP levels under conditions relevant for infection, because OccAB1/OprD was previously identified as abundant *in vitro* (28) while our analysis shows it is not detectable *in vivo*. Of the other OccAB channels, only OccAB2 (VanP) and OccAB4 (BenP) are detectable, but both are present in low copy numbers. In conclusion, our analysis identifies DcaP and possibly Omp33-36 as hijack targets for future antibacterials. Because predictions point to DcaP having a larger barrel than Omp33-36 (16 and 14 strands respectively) we opted to focus on DcaP.

### The crystal structure of DcaP shows a trimeric 16-stranded β-barrel

Structure predictions suggest that the first ~60 residues of mature DcaP are in the periplasmic space. We therefore expressed full-length (DcaP_fl_), a truncated version lacking 40 residues at the N-terminus of the mature protein (DcaP_DN40_) and a barrel-only protein lacking the first 59 residues (DcaP_Trunc_) in *E. coli* and obtained reasonable yields for OM expression (~3 mg from 12 litres of culture; Methods). Fig. S3 shows that boiled and non-boiled SDS-PAGE samples have different mobilities, demonstrating that DcaP proteins are heat-modifiable and form stable beta-barrels. However, for boiled DcaP_fl_ and DcaP_DN40_ constructs, two major SDS-bands are visible at around 45 kDa. Mass spectrometry analysis identified the upper band as having the theoretical mass of both constructs. The lower molecular weight band has the same size for both constructs and starts with amino acid 48, indicating proteolytic degradation. All DcaP constructs elute as trimers from size exclusion chromatography (SEC) columns. The purified proteins, containing a mixture of the full-length and the degraded protein species, were purified by Nickel-affinity chromatography and SEC, and crystallized in the presence of 0.4% C_8_E_4_. Crystal trials were initially set up for the full-length protein, but even after optimization crystal diffracted only to ~ 6 Å. By contrast, DcaP_DN40_ crystals diffracted much better (up to ~2.2 Å). However, structure solution using molecular replacement failed, due to the absence of a good homology model. Therefore, the structure was solved by single anomalous dispersion (SAD) phasing using seleno-methionine (SeMet). For this, we mutated two leucine residues (L280 and L282) to methionines in DcaP_DN40_ for increased phasing power (DcaP contains only 2 endogenous methionines), expressed this construct in minimal media containing SeMet (Methods; Table S1), and purified the substituted protein in the same way as the native species.

The first 19 amino acids of DcaP_DN40_ are not defined in the electron density and the model starts therefore with Leu59 (numbering for the mature protein, starting with ^1^ATSD). In contrast to other solved crystal structures of OM proteins from *P. aeruginosa* and *A. baumannii* which all crystallize as monomers (13, 15,35), DcaP crystallizes as a trimer containing three barrels with 16 β-strands each, thereby marking DcaP as the first trimeric OM protein from *A. baumannii* (Fig. 2 A,B). A DALI search yielded several trimeric OMPs as closest homologs *(Providencia stuartii* OmpPst2 and *E. coli* OmpC porins, as well as *E. coli* PhoE), with Cα r.m.s.d’s of ~3 Å and Z-scores between 24 and 25. In many porins, the N-and the C-terminal residues form a salt bridge within the last β-strand. In DcaP however, the N-terminus is in the periplasmic space, and the C-terminal amino acid F405 makes a hydrogen bond via the carboxylic group to the sidechain hydroxyl of S65 in a neighbouring barrel.

**Figure 2.**
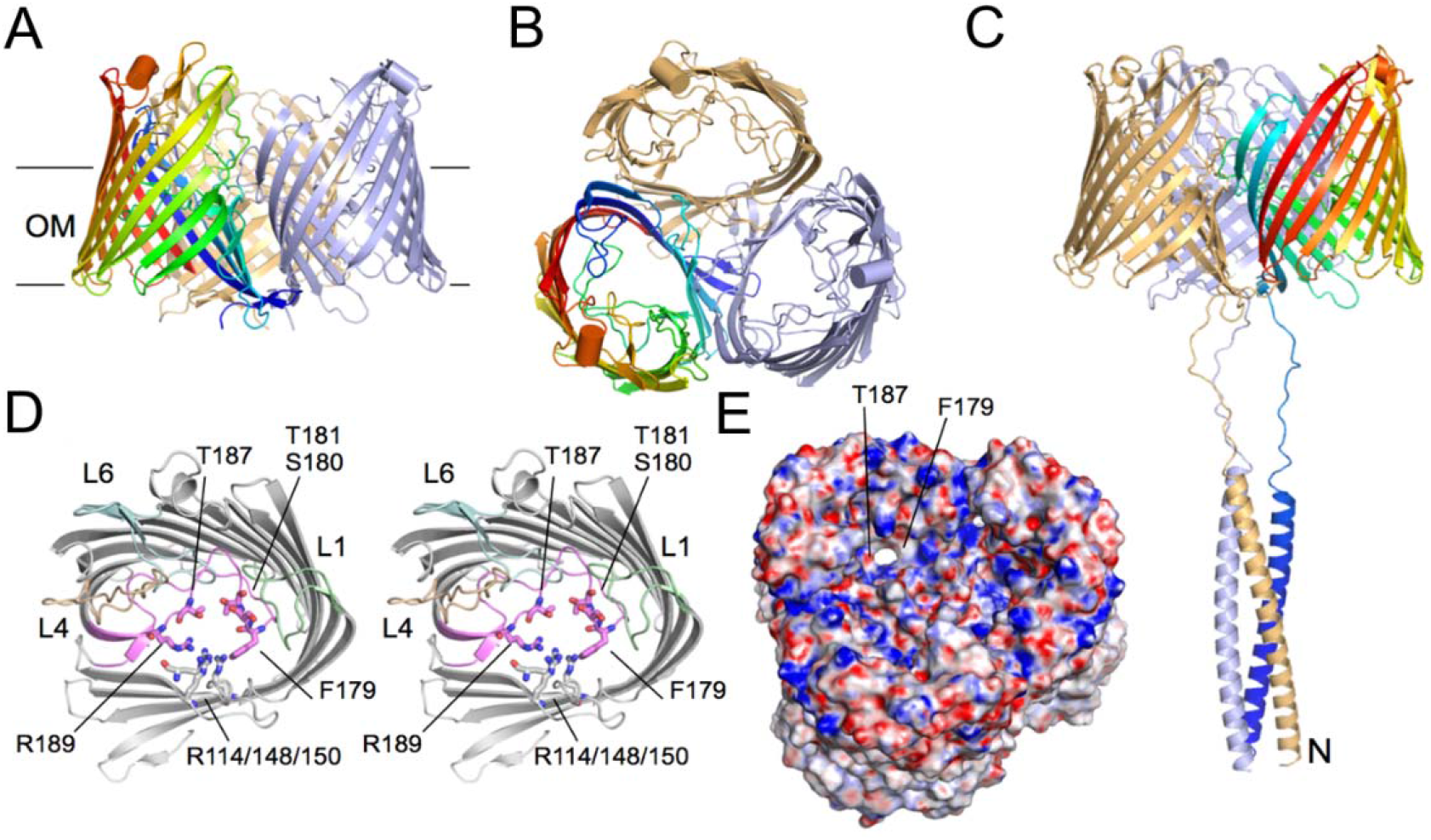
DcaP forms a trimeric 16-stranded β-barrel with a periplasmic extension. A, B Cartoon representations of the trimeric DcaP crystal structure viewed from the OM plane (A) and from the extracellular side (B). C, Side view of DcaP with the modelled N-terminal 59 residues added in cartoon representation. D, Stereo view of the constriction region, with loop L3 coloured pink. The residues lining the constriction region are labelled. E, Electrostatics surface view of the DcaP trimer from the outside of the cell, contoured from −20 to 20 ke/T. The view is at an angle to show one pore of the trimer clearly. The structural figures were made using Pymol (29).

Sequence similarity searches revealed the presence of a coiled-coil in the N-terminus, similar to the sucrose-specific channel ScrY in which first 42 amino acids of the mature sequence form a coiled-coil domain (36). To obtain a model for the entire DcaP structure, we modelled the first 60 amino acids of the mature protein, residing in the periplasmic space (Methods). The full-length predicted structure comprises a 49-residue coiled region that is connected to the barrel via a 10-residue disordered linker (Fig. 1C). An atomistic molecular dynamics simulation of the full-length DcaP structure revealed that the N-terminal domain undergoes rapid movements, which must be due to the disordered linker (Fig. S5 A). However, the trimeric coiled-coil domain itself remained very stable throughout the simulation, and extends up to ~100 Å into the periplasmic space. Interestingly, almost one-third of the residues are glutamines, with a Gln-rich stretch between residues 20-40. Like porins and the substrate-specific Occ proteins from *A. baumannii* and *P. aeruginosa* (15, 31–33,35), DcaP contains several arginine residues in the wall of the barrel (Arg114/148/150/189), which are all located on one side of the constriction region (CR; Fig. 1D). On the other side of the constriction, porins have electronegative groups and negatively charged residues, and this creates a strong electric field across the CR that is crucial for substrate translocation (37). This typical configuration is absent in DcaP, with a phenylalanine (Phe179) and two threonine residues (Thr181 and Thr187) opposite the arginines. While those residues and several pore-lining backbone carbonyls (Fig. 1D) provide electronegative groups, the overall contribution of the positive charges in the CR of DcaP is much higher than in porins, and this may generate specificity for negatively charged small molecules. Similar specificity might also be present in other related *Acinetobacter spp*. as the arginine residues located in the CR are conserved among various other aligned DcaP sequences (Fig. S1).

### The N-terminal coiled-coil domain stabilizes the DcaP oligomer

Multi-channel reconstitution of the DcaP_fl_ protein in planar lipid bilayers reveals stepwise insertions with variable conductance states (Fig. 3A). The corresponding histogram distribution reveals a major peak around 1500 pS, whereas minor peaks are observed around 500 and 1000 pS that might correspond to monomeric and dimeric channels (Fig. 3B). The high-resolution crystal structure suggests that the relatively unstructured N-terminal domain might cause transient blockages of the channel by interacting with the periplasmic side of the pore, and therefore we also tested two truncated versions of DcaP; one containing only the β-barrel domain (DcaP_Trunc_) and another lacking the first 40 amino acids of the N-terminus (DcaP_DN40_). The corresponding histograms overlapped with that of wild type DcaP_fl_ (Fig. 3B) and the major conductance peaks have shifted in the shorter variants towards the conductance value for a monomer (~500 pS), suggesting that the N-terminus plays a role in stabilization of the trimer. To support this hypothesis, we estimated the interaction energy of one monomer with the other two monomers from the unbiased MD simulations for DcaP_fl_ and the barrel-only DcaP_Trunc_ variant, and compared the energies with those of OmpC, known to form very stable trimers (38, 39). For the DcaP_Trunc_ variant, the stability of the barrel is lower compared to OmpC (Fig. S5B). By contrast, the full-length protein is more stable than OmpC, suggesting that the N-terminal coiled-coil domain provides additional oligomer stability (Figs. S4 and S5).

**Figure 3.**
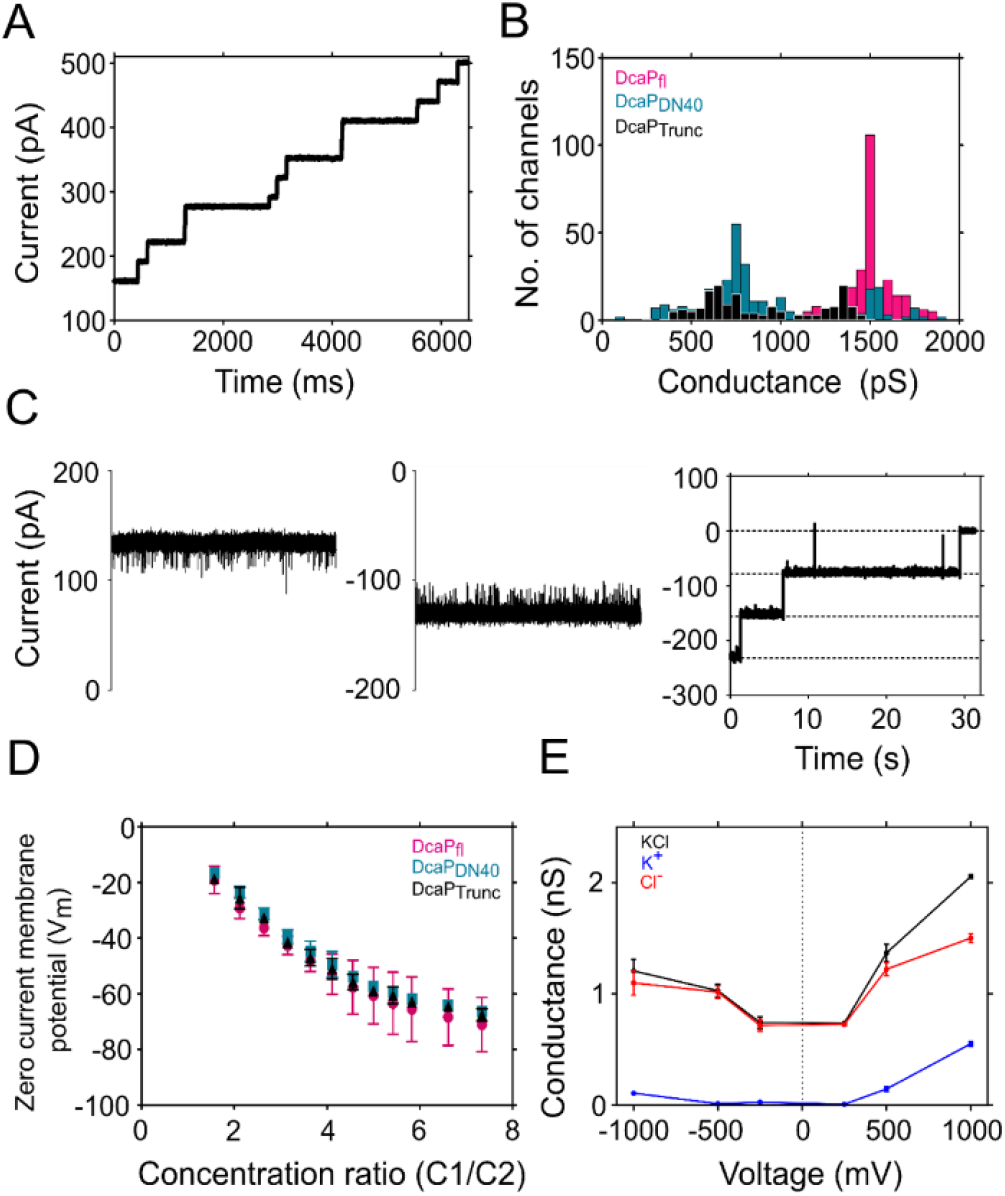
Characterization of DcaP by electrophysiology. A, Typical porin reconstitution into planar lipid bilayer at 1 M KCl 10 mM HEPES pH 7 and 50 mV applied voltage. Individual channel insertion is observed by sudden current jump. B, Plot of the number of channel insertion events vs. channel conductance. A bimodal current distribution is observed with maxima identified as mono or trimer conductance. C, Representative single channel ion current trace of DcaP_fl_ at +100 (left panel) and −100 mV (middle panel) in 1 M KCl 10 mM HEPES pH 7. Right panel, typical voltage-induced channel closure for DcaP_fl_ at high external applied voltage (−175 mV), showing sequential three-step closures for each monomer. D, Zero-current membrane potential measurements for all three DcaP variants, demonstrating specificity for anions. E, Conductance estimates for DcaP_Trunc_ obtained from applied field MD simulations in 1 M KCl with respect to the applied external voltages. The conductance values for the individual ionic species are shown as well.

We next moved to single channel experiments, in which DcaP_fl_ displayed a conductance of around 1.3 nS in 1M KCl (Fig. 3C). We observed trimeric gating behaviour of the channel at applied voltages above 150 mV, confirming that DcaP_fl_ is trimeric (Fig. 3C). For the truncated variants we observed insertions corresponding to monomeric, dimeric and trimeric channels, which is consistent with the behaviour observed in multi-channel experiments. In the experiments described below we have only considered trimeric channels, and have determined current (I) - voltage (V) relationships for all variants (Fig. S6A). The I - V curves are linear and overlap for the three proteins, suggesting that (i) DcaP forms open channels without voltage-induced gating by extracellular loops and (ii) that the N-terminal periplasmic domain does not contribute to the channel conductance.

### DcaP is strongly anion-selective

We next performed selectivity experiments to determine the ion selectivity of DcaP (33). Here, zero-current membrane potentials (40, 41) are determined over a range of concentration ratios between both sides of the membrane. For all DcaP variants we obtained high negative potentials (Fig. 3D), indicating that the channel has a strong preference for anions over cations. The estimated permeation ratio of 1:10,000 for K^+^: Cl^-^ suggests that DcaP is almost exclusively selective for anions. This correlates very well with the abundance of positively charged residues in the CR (Figs. 2D,E) and with the earlier genetic studies (18, 42), hinting at dicarboxylic acids as probable substrates. To complement the experimental observations, we performed applied field simulations to estimate the conductance and selectivity of DcaP_Trunc_ (Fig. 3E). The calculated conductance values are in good agreement with the experimental value for high voltages, though discrepancies remain at low voltages. The calculation of the separate conductance values for only potassium or only chloride ions confirm the electrophysiology results that identify the chloride ions as the major conducted species (Fig. 3E).

### Phthalates are transport substrates for DcaP

The DcaP crystal structure, together with the electrophysiology experiments, suggests that negatively charged compounds containing hydrophobic moieties might be potential transport substrates of DcaP. Phthalates meet these criteria, and we tested their interaction with DcaP using electrophysiology. We also tested succinic acid, a linear dicarboxylic acid. In addition, arginine was tested as an example of a net positively charged compound. Structures of the relevant substrates are shown in Fig. S13. Typical interactions of DcaP with substrates are shown in Fig. 4 and Fig. S7. In contrast to discrete ion current blockages observed in other channels resulting from strong substrate interaction in the constriction region (40,43–47), we instead observed a decrease in current upon addition of substrate. The current decrease was dependent on the concentration and type of substrate, with the largest current decrease observed with *o*-phthalic acid. Similar interactions but with smaller current decreases were observed with *m*-phthalic acid and *p*-phthalic acid (Fig. 4). Very small current decreases were observed with succinic acid. Arginine showed no interaction with the channel (Fig. 4E and Fig. S7).

**Figure 4.**
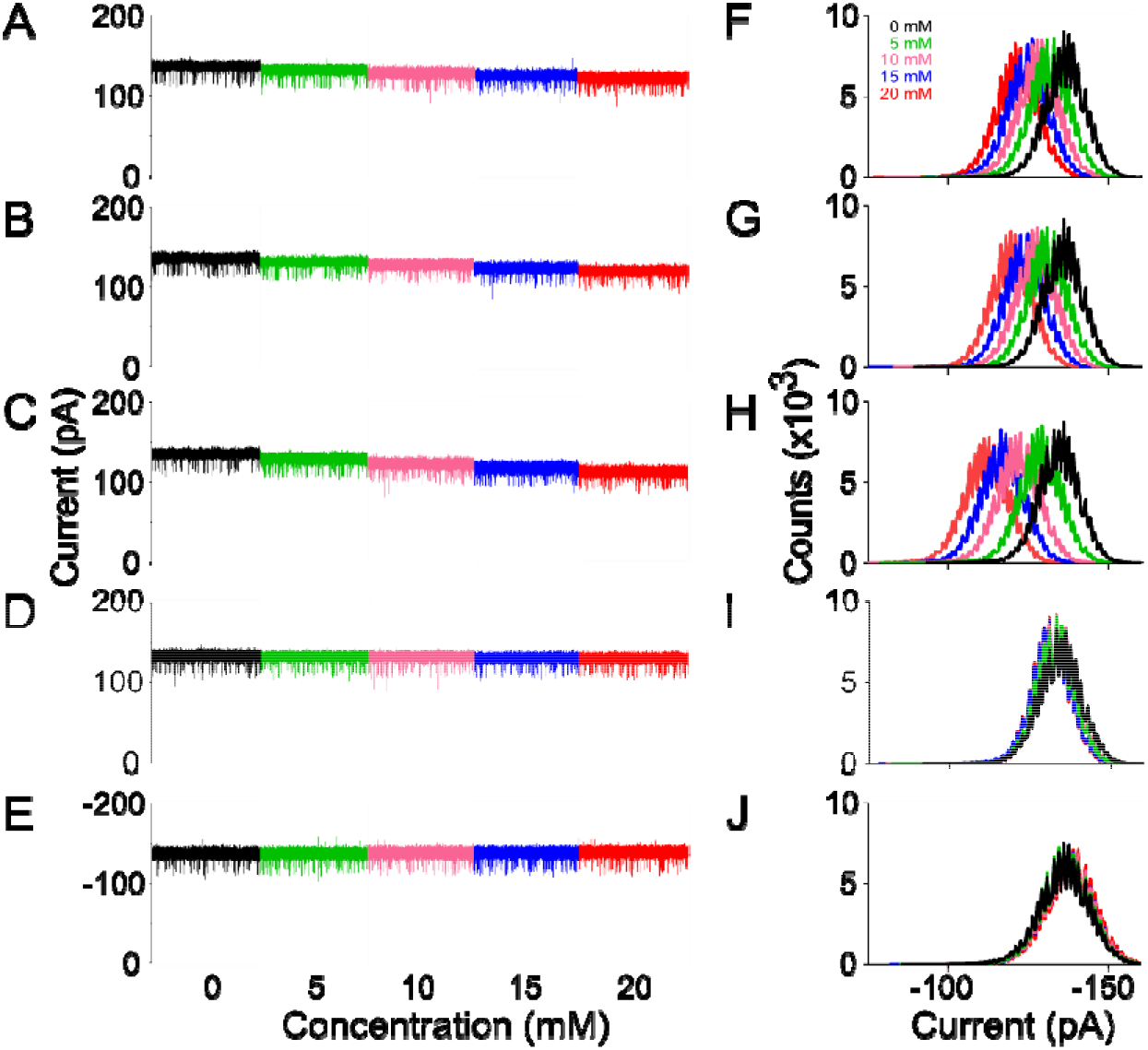
Phthalic acids are likely substrates for DcaP. A-J, Ion current traces and corresponding histograms showing interaction of DcaP with *o*-phthalic acid (A,F), *m*-phthalic acid (B,G), *p*-phthalic acid (C,H), succinic acid and (D,I) and arginine (E,J) at +100 mV applied voltage. All substrates were added on the cis side at 5-20 mM concentrations. The aqueous solution contained 1 M KCl and 10 mM HEPES pH 7.

In addition to the nature of the compound, the current decreases were also found to be dependent on the side of addition of the substrate. For most OM proteins after bilayer insertion, the extracellular side of the channel remains on the side where the protein was added (cis) (48). The reason for this is that the extracellular loops are mostly long and hydrophilic, making it unfavourable for those to cross the membrane spontaneously. By contrast, periplasmic turns tend to be short, making the energetic barrier for membrane translocation lower. From the concentration-dependent decreases, we evaluated the binding kinetics using Langmuir isotherms and found that *o*-phthalic acid binds with a binding affinity of around 9 mM from the cis side (extracellular side) and 25 mM from trans side (periplasmic side) (Fig. S8). For all three phthalates, binding affinities are similar from the extracellular side but differed from the periplasmic side. For *o*-phthalic acid, the affinity is higher from the periplasmic side as compared to the extracellular side whereas for *m*-phthalic acid and *p*-phthalic acid the extracellular side affinity is higher. There was no significant difference in substrate interaction between the full-length and barrel-only protein, confirming that the N-terminus does not play a role in substrate interaction (Fig. S7C, D).

To obtain a molecular picture of substrate permeation, applied field simulations were performed at −1 V and −0.5 V for all three cyclic phthalate isoforms, succinic acid and arginine (at + 0.5 V). We observed permeation for all negatively charged substrates at both applied voltages (Fig. S9). By contrast, even for high voltages we did not observe any permeation for arginine. Compared to the phthalic acids, the permeation of succinic acid is very fast and likely the reason why only very small current decreases were observed in single channel electrophysiology experiments (Fig. 4). As expected, the residence time for all molecules was increased in the CR (z= −7 to 4 Å) at 0.5 V compared to 1 V, indicating a significant affinity of the CR for these molecules. Moreover, all permeating substrates adopt very specific orientations during translocation, which is very similar at both applied voltages, (Fig. S10) and representative orientations of all four substrates are shown in Fig. 5. In all cases, the interaction is as expected from the CR structure, with the carboxyl groups interacting with the arginine side chains whereas the aromatic ring of the phthalic acids is oriented towards the more hydrophobic part of the CR.

**Figure 5.**
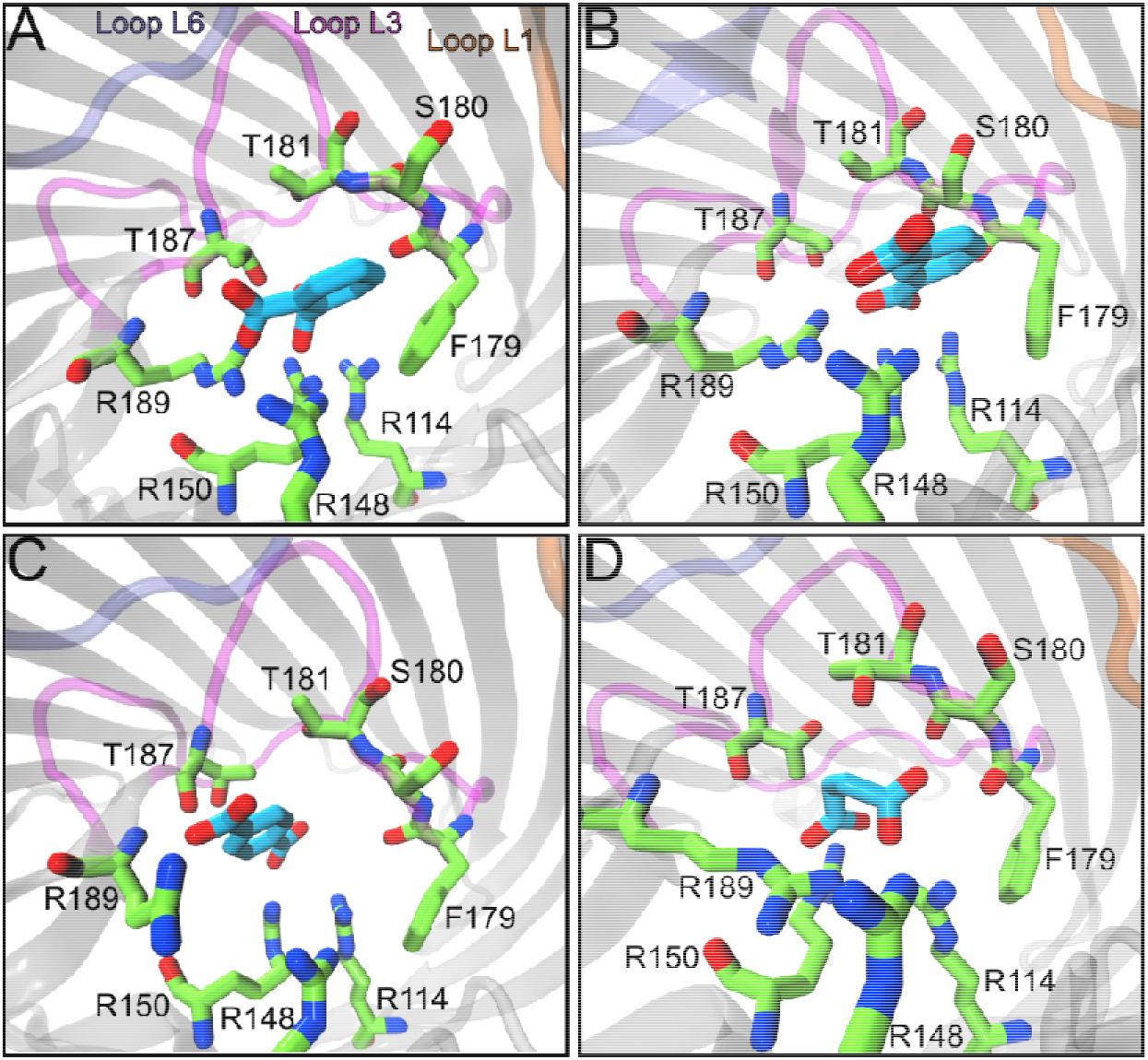
Substrate translocation via DcaP explored by applied field MD simulations. Representative conformations of the substrates *o*-phthalic acid (A), *m*-phthalic acid (B), *p*-phthalic acid (C) and succinic acid (D), near the CR of the channel. The substrates and interacting residues in the constriction region are shown in stick representation. Carbon atoms are depicted in cyan and green for the substrates and amino acids, respectively. Oxygen and nitrogen atoms are shown in red and blue, respectively.

To evaluate the potential relevance of DcaP for antibiotic uptake, we tested sulbactam, ticarcillin, ampicillin, and piperacillin because of their clinical relevance for treating *A. baumannii* infections. We observed interactions with all tested antibiotics. With sulbactam, we observed interactions like phthalates, *i.e*. a decrease in overall current without discrete current blockages, whereas with tazobactam, ticarcillin and ampicillin we observed both current decreases as well as discrete blockages (Fig. 6). However, with piperacillin no current decrease was observed except for blockages. Furthermore, we observed occasional long blockages up to several milliseconds, likely resulting from the obstruction of channel with a non-permeating antibiotic molecule.

**Figure 6.**
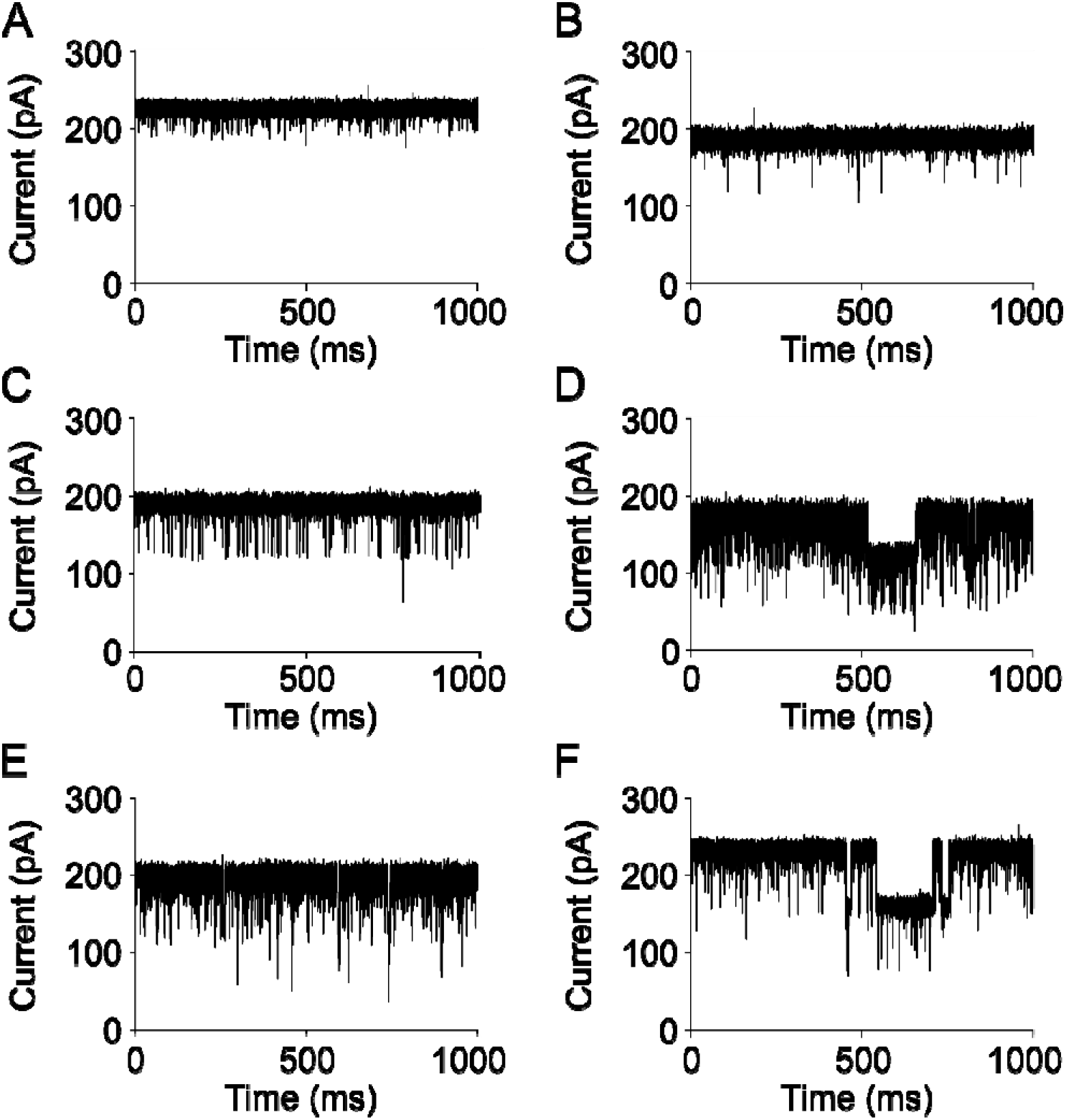
DcaP interacts with several antibiotics. Single channel current traces of DcaP_Trunc_ in the absence (A) and presence of 10 mM of (B) sulbactam, (C) tazobactam, (D) ticarcillin, (E) ampicillin and (F) piperacillin at +175 mV applied voltage. Antibiotic was added to the cis side of the membrane. The aqueous solution was buffered with 1 M KCl 10 mM HEPES pH 7.

A known problem in standard electrophysiology experiments is the inability to distinguish substrate binding/unbinding from translocation, since in both cases current decreases and/or blockages are observed. To obtain a better insight into possible antibiotic translocation we next performed applied field simulations as well as reversal potential measurements for one antibiotic, sulbactam. We selected sulbactam because of its relatively small size (233 Da), favouring translocation, as well as because of its similarity to the identified phthalic acid substrates (Fig. S13). In the applied field simulations, sulbactam tends to approach the CR quickly but remains on either side of the CR during the simulated timescale, indicating a significant affinity for the CR (Fig. S11), most likely due to the presence of the carboxyl and sulphone groups. We observed one permeation event from the periplasmic side of the pore, which suggests that DcaP is a probable permeation route for sulbactam. For a better molecular picture, we reconstructed the 2D free energy surface from metadynamics simulations and estimated the lowest energy path along the surface (Fig. 7). No significant interactions of sulbactam were observed away from the CR in both extracellular and periplasmic vestibules. By contrast, the presence of deep energy minima on the extracellular side of the CR (z ~ −5 Å) suggests a strong affinity of the molecule for channel residues (Fig. 7C). From these minima, the molecule can follow two permeation pathways to reach the minima located on the periplasmic side, with almost identical barriers of ~8 kcal/mol. However, the permeation barrier can be overcome more easily for permeations from the periplasmic side due to the intermediate steps with maximum step heights of ~4 or ~6 kcal/mol for path A and B, respectively. During translocation events from the extracellular side, an energetic barrier of ~8 kcal/mol must be overcome in one step. These findings support the directionality of the permeation event observed in the applied field simulations. The relatively low barriers are a strong indicator of the involvement of DcaP in sulbactam permeation.

**Figure 7.**
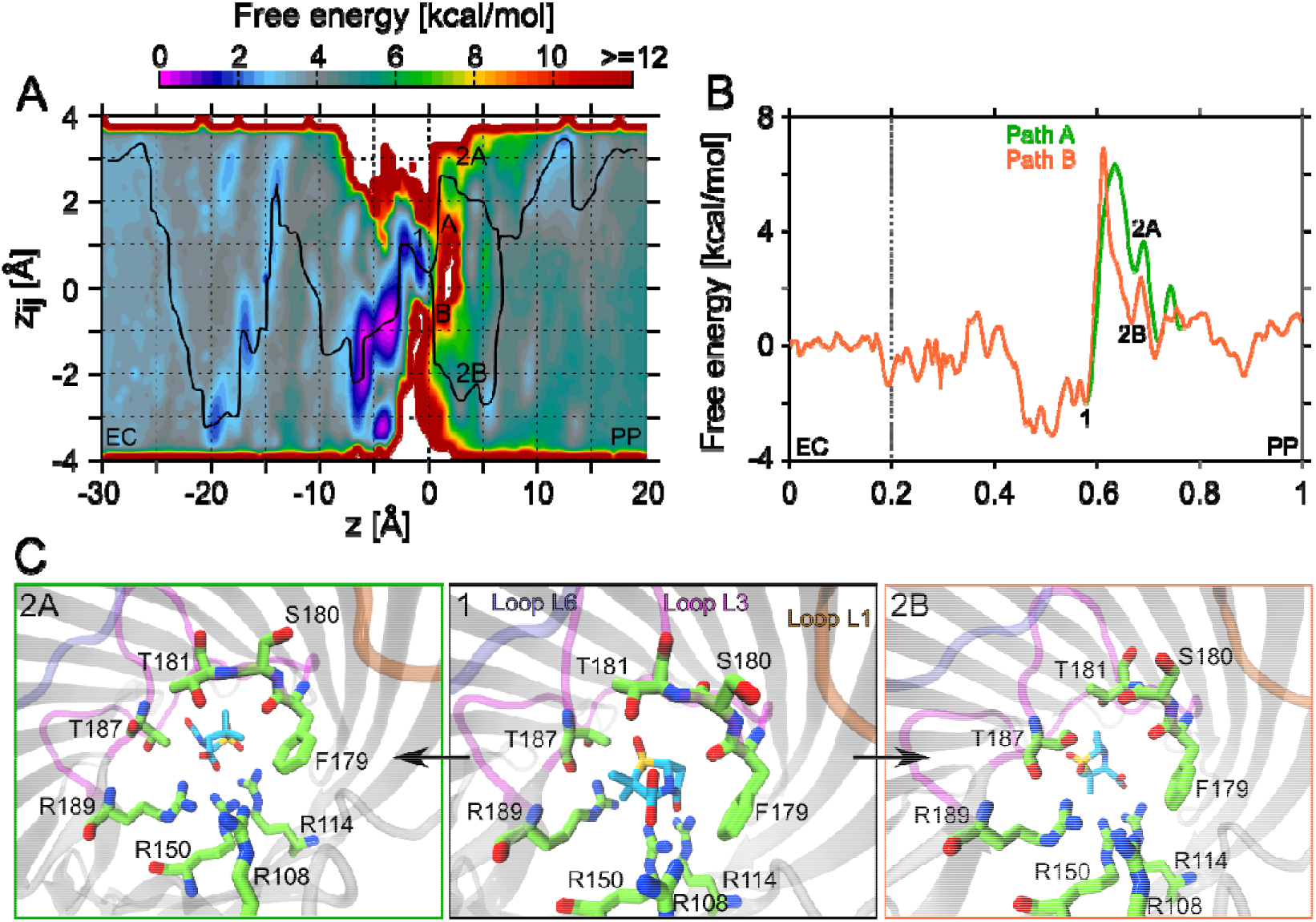
Interaction of sulbactam with DcaP. A, Reweighted free energy surface as a function of the collective variables *z* and *z_ij_* shown for sulbactam transport via DcaP. The two lowest-energy pathways are depicted as black lines. Important metastable states found in the constriction region are labelled. EC and PP denote the extracellular and periplasmic side, respectively. B, Free energy along the lowest-energy pathways for sulbactam. The curvilinear lowest-energy paths in (A) are parametrized by a variable s with s=0 and s=1 corresponding to the sulbactam being at the extracellular and the periplasmic side of the pore, respectively. The free energies are assumed to vanish at the extracellular side. C, Representative conformations of sulbactam and DcaP CR residues in free energy minima found before the major transition barriers from either side of the CR (1, 2A and 2B) are shown in stick representation. A similar colour scheme was used as in Fig. 5.

Finally, we used reversal potential measurements to directly detect translocation. Application of a concentration gradient of 25 mM Na-sulbactam creates a potential of around −14 mV (Fig. S12). At this reversal potential, the concentration-driven flux of sulbactam” balances the electric field-driven movement of Cl^-^ anion. According to the Goldman-Katz equation (49,50), the potential corresponds to a permeability ratio of K+: Cl^-^: sulbactam of^-^ 1:5000:1000. Extrapolation of the chloride conductance (1.2 nS measured from 1 M to 10 mM KCl) gives a chloride ion current of I = (12 × 10^−12^ S) × (14×10^−3^ V) = 1.6 × 10^−13^ A = ~10^6^ ions/s. This ion flow will be balanced by a 25 mM sulbactam gradient. With typical gradients of ~1 μM occurring in MIC assays, the sulbactam flux would correspond to ~ 4 - 8 molecules/s.

## Discussion

The low permeability of the OM plays an important role in both intrinsic and acquired antibiotic resistance of Gram-negative bacteria. This is especially true for pathogens like *P. aeruginosa, A. baumannii* and *B. pseudomallei*, which do not have abundant, permanently open OmpF and OmpC orthologs that mediate uptake of many antibiotics in Enterobacteria (45, 51). For such organisms, it is therefore crucial to (i) identify which OM channels are abundantly expressed during infection *in vivo* (as opposed to abundance in standard laboratory media), followed by (ii) structural and functional characterization of those channels to identify transport substrates. This knowledge can then be used to guide the design of antibiotic scaffolds that hijack the channel of interest for cellular entry. In this study, we have applied this approach to *A. baumannii*, an ESKAPE pathogen (52, 53) and recently identified by the WHO as a multidrug-resistant pathogen for which a critical need for novel antibiotics exists (54). As expected from the low permeability OM of this organism, most known and predicted diffusion channels are expressed at very modest levels (< 1000 copies/cell) when compared to the combined ~10^5^ OmpF and OmpC channels in *E. coli*. Our analysis shows that DcaP is the most highly-expressed diffusion channel in infected rodent lungs in *A. baumannii*. This identifies DcaP as a channel of outstanding interest for targeting by future antibacterials.

We have not tested whether clinically relevant antibiotics against *A. baumannii* require DcaP for cellular entry. In principle this could be done, for example, by comparing MICs of wild type and *dcaP* knockout strains. However, we consider it unlikely that DcaP facilitates entry of existing antibiotics (at least to the extent of lowering MICs) because these have not been designed to permeate efficiently, neither via DcaP nor via any other OM channel of *A. baumannii*. The fact that no mutations in *dcaP* have been reported in drug-resistant *A. baumannii* strains supports this notion and underscores the potential of DcaP as a novel drug delivery channel. A somewhat similar argument can be made for the *in vivo* verification of DcaP small molecule substrates that we have identified by electrophysiology *in vitro*. A *dcaP* knockout would only show growth defects on phtalates if DcaP is the only entryway for these compounds, which, given the versatility of bacteria, is unlikely. Indeed, we have demonstrated previously that OccAB3 and OccAB4 (BenP) are both anion-specific (15), and their potential upregulation in a *dcaP* knockout could mask dependence on DcaP. However, such redundancy would not diminish the potential of exploiting DcaP as a drug entry channel.

Interestingly, DcaP has clear structural similarities with classical 16-stranded trimeric porins such as OmpC and PhoE and is distinct from the other structurally characterized, monomeric *A. baumannii* OMPs, *i.e*. the 18-stranded OccAB channels (15) and the eight-stranded CarO protein (13). Another unusual property of DcaP is the long N-terminal periplasmic extension. So far, very few OM proteins are known with such N-terminal extensions, with examples being the sucrose channel ScrY and the putative amyloid secretion protein FapF from *Pseudomonas aeruginosa* (36, 55). The role of these N-termini in passive diffusion channels remains somewhat unclear. For the 18-stranded sucrose channel ScrY, the 70 amino acids at the N-terminus have been proposed to form a coiled-coil periplasmic binding protein which acts as a sink for the incoming substrate and might facilitate subsequent transport across the inner membrane. (36, 56) For FapF, the situation is a bit more complicated. Like in DcaP, the N-terminal ~80 residue domain is predicted to form a coiled-coil and needed removal before well-diffracting crystals could be obtained. In the FapF crystal, a helical segment that directly precedes the barrel plugs the channel from the periplasmic side. Since single-channel experiments suggested that the full-length channel is open, the helical plug domain was proposed to be involved in gating of the channel during amyloid export. Intriguingly, a protease present in the *fap* operon could, via cleavage of the N-terminal domain, result in plugging of the barrel after amyloid export (55). For DcaP, the electrophysiology experiments suggest that the N-terminus does not play a role in substrate transport and is not involved in channel gating. Since DcaP is not part of an operon and, based on its structure, most likely functions as a classical diffusion import channel, we hypothesize that the ~60 N-terminal amino acids of DcaP may have a role in maintaining structural integrity of the trimer and might play a ScrY-like role as a periplasmic binding protein guiding the substrate to the yet unidentified inner membrane transporter. Further biochemical and modelling approaches are required to elucidate the exact role of the DcaP N-terminus.

Much is known about the presence of catabolic island clusters in *Acinetobacter* species involved in the degradation of different compounds. Examples are the *dca-pca-qui-pob* operons which are independently transcribed and allow growth on dicarboxylic acids *(dca)*, protocatechuate *(pca)*, quinate *(qui)*, and p-hydroxybenzoate *(pob)* (18, 42, 57). By contrast, virtually nothing is known about how such compounds are taken up by the cell, a prerequisite for their degradation. Using electrophysiology and computational techniques, we have identified DcaP as an uptake channel for dicarboxylic acids, including phthalates. In terms of structure, DcaP is similar to phosphoporin PhoE, which has a 3-4-fold preference for anions over cations (41). Due to the strong preference of DcaP for anions, DcaP exhibits functional similarity with OprP of *P. aeruginosa*, which has a preference ratio of 100 for anions over cations and is also involved in phosphate uptake (58). Interestingly, several environmental strains of *Acinetobacter* that contain DcaP can degrade phthalates and utilize them as carbon sources (59), suggesting phthalates are physiologically relevant transport substrates. The strong selectivity of DcaP for negatively charged substrates could have interesting implications for narrow-spectrum antibacterials. The porins OmpF and OmpC of Enterobacteria are invariably cation-specific (41), and the presence of a primary amine in the substrate is known to facilitate uptake (60). This is unlikely to be the case for DcaP, and the molecular rules for OM permeation in *A. baumannii* might be quite different from *e.g. K. pneumoniae*. This might be exploited via development of narrow-spectrum drugs, which, in combination with rapid diagnostic testing, would present many advantages compared to the indiscriminate use of broad-spectrum antibacterials.

Understanding how antibiotics and other small molecules pass through OM channels should enable the rational design of novel antibiotics with superior permeation properties. We propose that hijacking OM channels that are highly expressed during infection for the delivery of antibiotics that resemble the natural substrates of that channel, presents an attractive and untested approach for developing new antibacterial leads. While the low molecular weight of diffusion channel substrates will make structure-based antibiotic design challenging, a proof-of-principle example is provided by imipenem, which has a relatively low molecular weight (300 Da) and adventitiously resembles the arginine substrate of the OccD1/OprD channel of *P. aeruginosa* (32, 61). Interestingly, removal of OccD1 increases the MICs for imipenem (62–64), demonstrating that this antibiotic indeed utilizes OccD1 for OM passage. Another example, illustrating that small pores do not necessarily prevent passage of large molecules, is provided by albicidin from *Xanthomonas albilineans*, which is an oligopeptide of non-natural amino acids and a potent inhibitor of DNA gyrase (65). Albicidin, despite its large size (842 Da), is thought to traverse the OM via the very small pore of the nucleoside channel Tsx (66, 67), indicating that relatively large molecules can pass through small OM channels.

With respect to existing antibiotics against *A. baumannii*, we have demonstrated that the β-lactamase inhibitor sulbactam translocates through DcaP at appreciable rates *in vitro* even at typical, shallow concentration gradients. It would be of interest to test *in vivo* whether, and to which extent, sulbactam accumulation in *A. baumannii* depends on DcaP. While sulbactam alone has weak activity against *A. baumannii*, its prospects as a useful antibiotic were recently revived via combination with ETX2154, a novel compound derived from avibactam (68). Sulbactam is negatively charged and superficially resembles the transport substrate phthalic acid, providing additional support for feasibility of the “hijacking” approach we describe here. We suggest that further, DcaP-guided modification of sulbactam and avibactam-like compounds guided by the DcaP channel might result in better-permeating compounds for *A. baumannii*. In conclusion, we suggest that the multi-disciplinary approach we describe here for *A. baumannii* DcaP can be applied to any OM channel and provides a starting point for the design of novel and better permeating drugs against multidrug resistant Gram-negative bacteria.

## Methods

### Animal infection models

*Intra-bronchial instillation model*: specific pathogen free (SPF) immunocompetent male Sprague-Dawley rats weighing 100 - 120 g or male CD-1 mice weighing 20 - 25 g were infected with an agar suspension containing approximately 10^7^ colony-forming units *Acinetobacter baumannii* ATCC 19606, deep into the lung via nonsurgical intra-tracheal intubation (69). In brief, animals were anesthetized with isoflurane (5%) and oxygen (1.5 L/min), infected via intra-bronchial instillation of (rats-100 μl) (mice-20 μl) molten agar suspension via intra-tracheal intubation, and then allowed to recover. At 24 h post infection, animals were euthanized and lungs were homogenized in 1 mL phosphate-buffered saline supplemented with 100 pg/mL tetracycline (PBS-Tet buffer). All procedures are in accordance with protocols approved by the GSK Institutional Animal Care and Use Committee (IACUC), and meet or exceed the standards of the American Association for the Accreditation of Laboratory Animal Care (AAALAC), the United States Department of Health and Human Services and all local and federal animal welfare laws.

### Sample workup for proteomics

The sample workup protocol was optimized to deplete host material while maintaining *A. baumannii* viability until lysis. All buffers and equipment were used at 0 to 4°C to minimize proteome changes during sample workup. The lung homogenate was filtered through a cell strainer and 2 mL of PBS-Tet buffer was added followed by vigorous vortexing for 30-60 s. After centrifugation at 500xg for 5 min, the supernatant was transferred to a fresh tube, and the pellet was extracted again with 1 mL PBS-Tet buffer. The supernatant was combined with the first supernatant and centrifuged again at 500xg for 5 min. The resulting supernatant was centrifuged one final time at 16’000 x g for 3 min. The pellet was resuspended in 1 mL 0.1% TritonX-100-Tet in ddH20, vortexed vigorously for 1 min, and centrifuged for 3 min at 16000xg. The supernatant was removed, and the pellet was stored at −80°C. Samples were thawed and sonicated for 2 × 20 s (1 s interval, 100% power). Proteins were alkylated with 10□mM iodoacetamide for 30□min in the dark at room temperature. Samples were diluted with 0.1M ammonium bicarbonate solution to a final concentration of 1% sodium deoxycholate before digestion with trypsin (Promega) at 37°C overnight (protein to trypsin ratio: 50:1). After digestion, the samples were supplemented with TFA to a final concentration of 0.5% and HCl to a final concentration of 50□mM. Precipitated sodium deoxycholate was removed by centrifugation at 4°C and 14’000□rpm for 15 min. Peptides in the supernatant were desalted on C18 reversed phase spin columns according to the manufacturer’s instructions (Macrospin, Harvard Apparatus), dried under vacuum, and stored at −80°C until further processing.

### Parallel reaction monitoring

Heavy proteotypic peptides (JPT Peptide Technologies GmbH) were chemically synthesized *A. baumannii* outer membrane proteins. Peptides were chosen dependent on their highest detection probability and their length ranged between 7 and 20 amino acids. Heavy proteotypic peptides were spiked into each sample as reference peptides at a concentration of 20 fmol of heavy reference peptides per 1 μg of total endogenous protein mass. For spectrum library generation, we generated parallel reaction-monitoring (PRM) assays (70) from a mixture containing 500 fmol of each reference peptide. The setup of the μRPLC-MS system was as described previously (71). Chromatographic separation of peptides was carried out using an EASY nano-LC 1000 system (Thermo Fisher Scientific) equipped with a heated RP-HPLC column (75 μm x 37 cm) packed in-house with 1.9 μm C18 resin (Reprosil-AQ Pur, Dr. Maisch). Peptides were separated using a linear gradient ranging from 97% solvent A (0.15% formic acid, 2% acetonitrile) and 3% solvent B (98% acetonitrile, 2% water, 0.15% formic acid) to 30% solvent B over 60 minutes at a flow rate of 200 nl/min. Mass spectrometry analysis was performed on Q-Exactive HF mass spectrometer equipped with a nanoelectrospray ion source (both Thermo Fisher Scientific). Each MS1 scan was followed by high-collision-dissociation (HCD) of the 10 most abundant precursor ions with dynamic exclusion for 20 seconds. Total cycle time was approximately 1 s. For MS1, 3e6 ions were accumulated in the Orbitrap cell over a maximum time of 100 ms and scanned at a resolution of 120,000 FWHM (at 200 m/z). MS2 scans were acquired at a target setting of 1e5 ions, accumulation time of 50 ms and a resolution of 30,000 FWHM (at 200 m/z). Singly charged ions and ions with unassigned charge state were excluded from triggering MS2 events. The normalized collision energy was set to 35%, the mass isolation window was set to 1.1 m/z and one microscan was acquired for each spectrum.

The acquired raw-files were converted to the mascot generic file (mgf) format using the msconvert tool (part of ProteoWizard, version 3.0.4624 (2013-6-3)). Converted files (mgf format) were searched by MASCOT (Matrix Sciences) against normal and reverse sequences (target decoy strategy) of the UniProt database of *Acinetobacter baumannii* strains ATCC 19606 and ATCC 17978, as well as commonly observed contaminants. The precursor ion tolerance was set to 20 ppm and fragment ion tolerance was set to 0.02 Da. Full tryptic specificity was required (cleavage after lysine or arginine residues unless followed by proline), three missed cleavages were allowed, carbamidomethylation of cysteins (+57 Da) was set as fixed modification and arginine (+10 Da), lysine (+8 Da) and oxidation of methionine (+16 Da) were set as variable modifications. For quantitative PRM experiments the resolution of the orbitrap was set to 30,000 FWHM (at 200 m/z) and the fill time was set to 50 ms to reach a target value of 1e6 ions. Ion isolation window was set to 0.7 Th (isolation width) and the first mass was fixed to 100 Th. Each condition was analyzed in biological triplicates. All raw-files were imported into Spectrodive (Biognosys AG) for protein and peptide quantification.

### Cloning, expression and purification of OM-expressed DcaP

DcaP was amplified from *Acinetobacter baumannii* (strain AB307-0294) genomic DNA (uniprot A0A0B9X9I7). In addition to the full length DcaP (DcaP_fl_), a truncated mutant missing the first N-terminal 40 amino acids was generated (DcaP_DN40_). Both constructs added a hexa-histidine tag to the C-terminus for purification by IMAC. For OM expression, the genes were cloned into an arabinose-inducible pB22-vector via *XbaI* and *XhoI*. The plasmids were transformed into *E. coli* C43(DE3) cells. After adding 0.1% arabinose at an OD600 of 0.6, cultures were incubated overnight at 20 °C and 150 rpm to induced expression. To solve the crystallographic phase problem with selenomethionine (SeMet)-phasing, two leucines close together within the OM region were mutated to methionines (L280M, L282M) to have four methionines in total. The cells were grown in LeMasters-Richards minimal media with glycerol as a carbon source (0.3% w/v) to OD_600_ ~ 0.6 at 37°C before SeMet in combination with lysine, phenylalanine, threonine, leucine, isoleucine and valine were added (72). Half an hour later the cells were induced with 0.5 % arabinose and the cultures were incubated for 6 hours at 30 °C and 150 rpm.

After cell disruption (Constant Sytems 0.75 kW operated at 20-23,000 psi), the membranes were harvested by centrifugation for 45 mins at 42,000 rpm (45 Ti rotor, Beckman), and resuspended in TBS buffer (20 mM Tris, 300 mM NaCl pH 8) with 3% Elugent (Calbiochem). After one hour incubation at room temperature, insoluble particles were removed by 45Ti centrifugation for 30 mins at 42,000 rpm and the supernatants were loaded onto a 10 ml nickel column. The column was washed with 10 column volumes (CV) TBS containing 0.2% lauryldimethylamine *N*-oxide (LDAO) and 25 mM imidazole. The proteins were eluted with 3 CV TBS containing 0.2% LDAO and 250 mM imidazole. Afterwards, the proteins were purified by two rounds of size-exclusion chromatography, first with a HiLoad 26/600 superdex 200 column (GE Healthcare) using 10 mM HEPES, 100 mM LiCl and 0.05% LDAO, pH 7.5, followed by a second gel filtration run with a HiLoad 16/600 superdex 200 (GE Healthcare) using 10 mM HEPES, 100 mM LiCl, 0.4% C_8_E_4_, pH 7.5. The purified proteins were concentrated to ~10 mg/ml and directly flash-frozen into liquid nitrogen.

### Crystallization and structure solution of DcaP

Initial crystallization trials for DcaP_fl_ and DcaP_DN40_ as well as their SeMet variants were performed at 295 K by sitting-drop vapour diffusion using the MemGold, MemGold2 and Morpheus screens from Molecular Dimensions with a Mosquito robot (TTP Labtech). The initial hits were optimized by manual fine-screening with larger drops by hanging drop vapour diffusion. DcaP_fl_ crystals were grown at 19°C by adding 1 μl as well as 1.5 μl of 10 mg/ml protein to 1 μl of reservoir solution, consisting of 26% PEG 400 and 0.1 M sodium citrate pH 5.5. However, the obtained thick hexagonal plates showed only poor and anisotropic diffraction patterns. For DcaP_DN40_, the crystallisation condition consisted of 8.5% MPD, 8.5% PEG 1000, 8.5% PEG 3350, 0.02 M sodium formate, 0.02 M sodium citrate, 0.02 M sodium oxamate, 0.02 M ammonium acetate, 0.02 M sodium potassium tartrate, 0.05 M MES, 0.05 M imidazole pH 6.5 (derived from the Morpheus screen). Crystals were directly flash-frozen in liquid nitrogen. A SAD data set was collected at the Diamond Light Source (DLS; Table S1), Didcot, UK. Data were integrated and scaled with XDS (73). Initial phasing and model building was done using AUTOSOL within PHENIX (74, 75). Further model building was performed using the program COOT (78). The protein model was refined with REFMAC.(69) The programs MolProbity (79) and PROCHECK (80) were used to evaluate the final model. PyMOL (29) was used for the visualization of the protein structure and for making figures.

### Electrophysiology

Conductance measurements were done for all DcaP protein variants using multichannel as well as single channel techniques in 1 M KCl 10 mM HEPES pH 7. Multichannel measurements were carried out as described elsewhere in greater detail (15). In brief, protein was added to the side (ground) of the chamber at 100 mV applied voltage. Histogram of the conductance *versus* the number of channels were plotted. Single channel measurements were done by using the Montal and Mueller technique (80). A ~100 μm aperture containing 25 μm thick Teflon film was partitioned between the two chambers of a Teflon cuvette. A solution containing 1 % hexadecane in hexane was used to paint the area around aperture to make the surface hydrophobic. A pair of Ag / AgCl electrodes immersed in the aqueous solution containing 1 M KCl 10 mM HEPES pH 7 were used to measure the electric current. 5 mg /ml DPhPC in pentane was used to form the membrane. Detergent-solubilized protein was added to the cis side of the membrane. One electrode was connected to the ground or cis side, with the other connected to the trans side and head stage of a Axopatch 200 B amplifier. Current traces were filtered by low pass Bessel filter at 10 kHz and sampled at 50,000 Hz. Data was analysed using Clampfit software (Axon Instruments, Foster City CA, USA) and plotted using Prism or Origin softwares.

Ion selectivity measurements were done as described elsewhere (41). Several hundred channels were reconstituted at low salt concentration of 0.1 M KCl 10 mM HEPES pH 7. After attaining saturation, salt concentration gradients were established by subsequent additions of 3 M KCl on the cis side and 0.1 M KCl on the trans side. Channel selectivity arising from asymmetric charge distribution in the CR leads to the preferential uptake of cations or anions, which gives rise to a potential value measured at the trans side (low dilution side) termed zero current membrane potentials. Negative potentials indicate preferential uptake of anions from cis side to the trans side.

Reversal potential measurements (49,50) were done to study the translocation of antibiotic sulbactam. In brief, channels were reconstituted in low salt such as 0.01 M KCl 1 mM HEPES pH 7. After attaining saturation driven by stable current value, asymmetric concentrations (25 mM, 50 mM) of sulbactam were added on the cis side of the chamber and resulting potentials values were noted measured on the trans side indicating the net transferred charges. A shift in the voltage (current) indicates the transferred charges. In this case, negative potentials indicate translocation of sulbactam as DcaP is a highly anion selective channel and the sulbactam counter (cat) ions cannot permeate.

### N-terminal structure prediction of DcaP

The first 41 N-terminal amino acid residues of DcaP are annotated to form a coiled-coil structure according to the uniprot entry A0A0B9X9I7. The submission of the full length sequence of DcaP to the PredictProtein webserver (81) predicts that the residues from 4 to 45 could from a coil. Moreover, visual inspection of the sequence suggests that coil formation might only be up to residue 49, due to the presence of several prolines (P50, P52, P56, P59) in the remaining 10 N-terminal residues that were not resolved in the crystal structure (Fig. S1). Therefore, the sequence of the first 49 amino acids was submitted to the CCBuilder webserver (82) to predict a trimeric coiled-coil structure. In total 36 structures were predicted by rotating the coils 10° relative to the long axis of the coiled-coil structure. To identify the most stable coiled-coil structure from the predicted ones, unbiased MD simulations were performed for all 36 structures by placing them into a water box. Each system was composed of ~ 31000 atoms. After energy minimization, the systems were equilibrated for 1 ns in NVT ensemble by maintaining the temperature at 300 K using a Nosé-Hoover thermostat and a time step of 1 fs. During this step, position restraints were applied on the protein atoms with a force constant of k = 1000 kJ mol^-1^ nm^-2^. The systems were further equilibrated in a NPT ensemble for 1 ns using a Parrinello-Rahman barostat by maintaining the pressure at 1 bar. The C atoms of the proteins were restrained during this step. Finally, unbiased MD simulations were performed for 50 ns in a NVT ensemble with a time step of 2 fs without applying any positional restraints. As shown in Fig. S5, the lowest root mean squared deviation (RMSD) was observed for coiled-coil structure #12. In addition, the calculated interactions between the coils was strongest for this structure and thereby showing the highest structural stability. Therefore, structure #12 was considered for further modeling. The full-length N-terminal structure prediction was performed in two steps. As mentioned earlier, the first 59 residues were not resolved in the crystal structure. Considering this crystal structure as a starting structure, the coordinates for residues from 50 to 59 were predicted using MODELLER version 9.11 (83). Subsequently, this structure and the equilibrated coiled-coil structure #12 were amalgamated using MODELLER to generate the full-length DcaP structure (Fig. 1C).

### Unbiased molecular dynamics simulations of WT DcaP

The full-length DcaP structure with the predicted N-terminal domain was inserted into a POPE lipid bilayer composed of 256 lipids. The system was further solvated using TIP3P water molecules and neutralized with 9 K^+^ ions. The resulting system consists of 280481 atoms in total. Following the minimization, the system was equilibrated in a NVT ensemble for 2 ns at a time step of 1 fs by maintaining the temperature at 300 K using the velocity rescaling (84) thermostat. Position restraints were applied to proteins and lipid head atoms during this step. Later on, the restraints were released from the lipid head groups and the system was further equilibrated in a NPT ensemble for 2 ns with a time step of 1 fs using the Nosé-Hoover thermostat (85) and the Parrinello-Rahman barostat (86) to maintain the pressure at 1 bar. The final step of equilibration of 5 ns was performed with a time step of 2 fs by only applying restraints of the backbone atoms of the protein. Furthermore, the system was simulated in a NVT ensemble for 200 ns without applying any restraints to understand the dynamics of the N-terminus.

### MD simulations of ions and substrates transport through DcaP channel

Applied field MD simulations (40, 87) were carried out to understand the permeation of ions (KCl), and substrates (*o*-phthalic acid, *m*-phthalic acids, *p*-phthalic acids, succinic acid, arginine). The truncated version of the DcaP structure (DcaP_Trunc_) was considered to reduce the computational expense. The system was built according to the above-mentioned system setup steps and a 1 M KCl solution was added. Following the minimization and equilibration steps, the simulations were performed at 0.25, 0.5 and 1 V at both voltage polarities for 250, 200 and 100 ns respectively, and repeated twice to estimate the IV-curve. Subsequently, 40 substrate or antibiotic molecules, equivalent to 80 mM concentration, were added to the same system. To understand the permeation from the extracellular (EC) to the periplasmic (PP) side, the systems including the substrates *o*-phthalic acid, *m*-phthalic acid, *p*-phthalic acid, or succinic acid were simulated for 200 ns and 500 ns at −1 V and −0.5 V, respectively. Due to the presence of the net positive charge on arginine, this applied field simulation was performed at +1 V for 200 ns. Due to the lack of translocation of arginine even at 1 V (see results section), simulations were not carried out at lower voltages. All simulations were performed using a time step of 2 fs.

### Dynamics and free energies of sulbactam translocation through DcaP

The applied field simulations for sulbactam were carried out with a time-step of 5 fs to achieve a longer timescale by using virtual hydrogen sites on the protein, lipids and antibiotic molecule (88, 89). The applied field simulations were carried out at both voltage polarities, i.e., at +1 and −1 V, for 1 μs to observe translocations from both sides of the channel. This set of simulations was carried out using a minimal number of ions only neutralizing the systems to be able to compare to the free energy calculations carried out using the well-tempered (90) and multiple walker metadynamics (91) simulations technique. The free energy calculations were performed as described elsewhere (88). Briefly, a total of 20 walkers were used with 12 and 8 walkers starting from the EC and PP side of the channel, respectively. During the first stage, the sampling of the sulbactam molecule was elevated along the channel axis by biasing the CV (collective variable) *z*, defined as the center of mass difference between the C_α_ atoms of the β-barrel of the respective monomer and the heavy atoms of the sulbactam molecule. The bias construction was carried out by depositing Gaussian hills with a width and height of 0.1 Å and 0.48 kcal/mol, respectively, at every 4 ps. The simulation for each walker was carried out for 200 ns leading to a total simulation time of 4 μs. Subsequently, a 2D free energy surface was estimated as function of CVs, *z* and *z_ij_*, where *z_ij_* basically defines the orientation of the molecule with respect to the channel axis using the Tiwary-Parrinello reweighting technique (92). The CV *z_ij_* represents the z-component of the interatomic vector connecting two atoms (see Fig. S10) which can be transformed to angles from 0 to 180 with respect to the z-axis. A similar CV was used to describe the orientation of substrates (Fig. S10). Finally, the minimum free energy pathways along the 2D free energy surface were estimated using the zero-temperature sting method (93).

All simulations were carried out using the GROMACS package version 5.1.2, (94) patched with PLUMED version 2.2.3. The CHARMM36 force field (95) was employed for all simulations. Moreover, the initial force field parameters for the substrates and antibiotic were taken from the CGenFF database (96) and optimized using the ffTK toolkit (97) as needed. The cut-off for the short-range electrostatics and the van der Waals interactions was set to 12 Å and the long-range electrostatics interactions were treated using the particle-mesh Ewald method (98) with a grid size of 1 Å. All bonds were constrained using the LINCS algorithm (99).

## Data Availability

The atomic coordinates and the associated structure factors have been deposited in the Protein Data Bank (http://www.pdbe.org) with accession code 6EUS. The authors declare that all other data supporting the findings of this study are available within the article and its Supplementary Information files, or are available from the authors upon request.

## Acknowledgements

We would like to thank the staff at beam line i03 of the Diamond Light Source UK for beam time (proposal mx9948) and assistance with data collection. The research leading to these results was conducted as part of the Translocation consortium (www.translocation.eu) and has received support from the Innovative Medicines Initiatives Joint Undertaking under Grant Agreement No. 115525, resources that are composed of financial contributions from the European Union’s seventh framework programme (FP7/2007-2013) and European Federation of Pharmaceutical Industries and Associations companies in-kind contribution. In addition, S.P.B and M.W acknowledge funding from Marie Skłodowska-Curie fellow within the ITN Translocation Network, project no.607694.

## Author contributions

M.Z and B.v.d.B purified, crystallized DcaP proteins and determined the crystal structure. S.P.B carried out and analysed electrophysiology measurements. J.D.P carried out and analysed molecular dynamics simulations. U.K. supervised the computational studies. J.H and J.W. prepared proteomics samples. C.S and S.S carried out in vivo proteomics supervised by D.B. B.v.d.B, D.B, U.K and M.W designed research. B.v.d.B and S.P.B wrote the paper, with input from all other authors.

## Additional information

Correspondence and requests for materials should be addressed to B.v.d.B, M.W and D.B.

---

**Fig S1.**
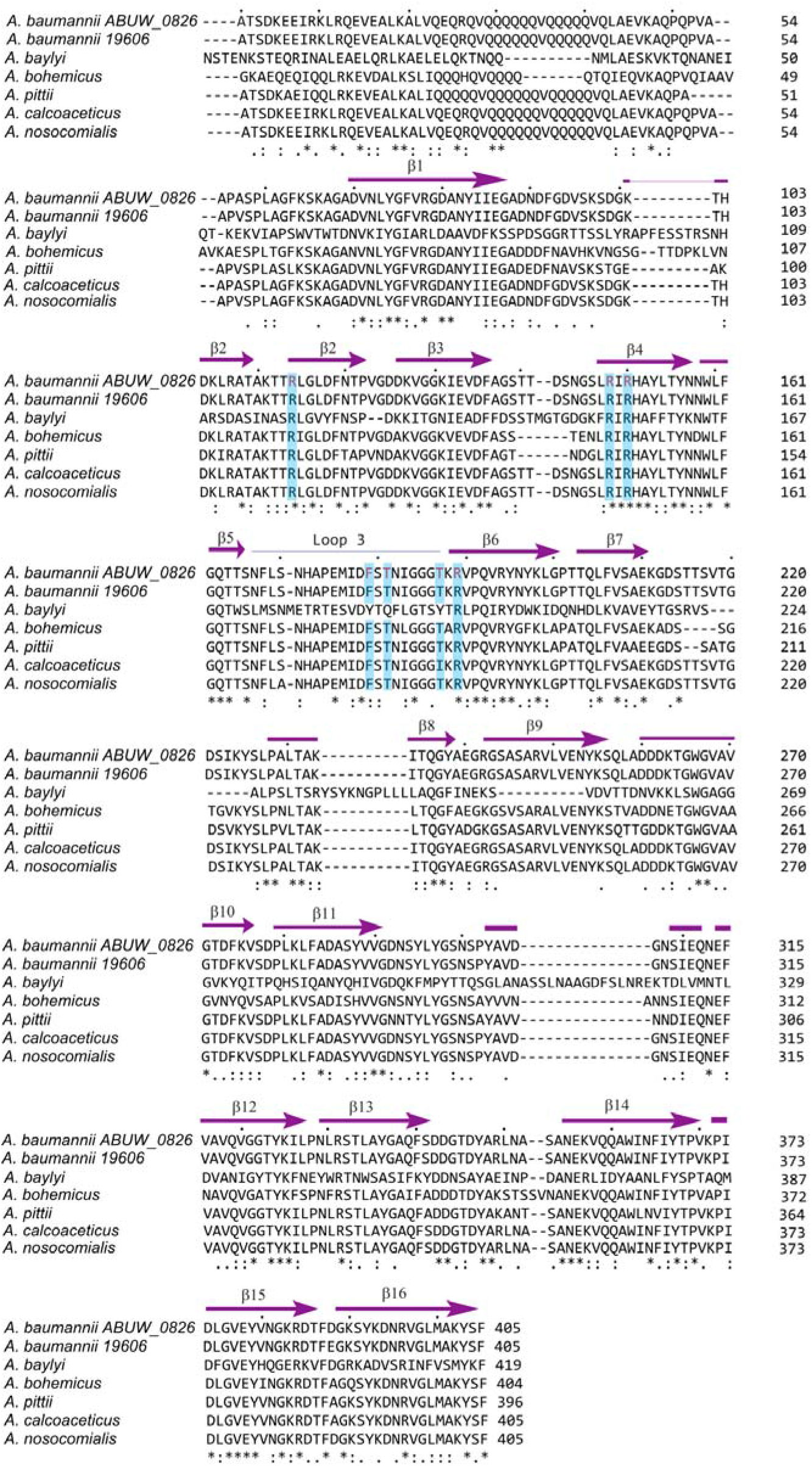
Sequence alignment of DcaP from *Acinetobacter* species with Uniprot IDs A0A0B9X9I7 (A. *baumannii ABUW _0826)*, N9LF65 (A. *baumannii strain 19606)*, Q937S8 (A. *baylyi (strain ATCC 33305 / BD413 / ADP1)*, N8Q9A1 *A. bohemicus ANC 3994)*, A0A0M3BW71 (A. *pittii)*, A0A1C4HM00 (A. *calcoaceticus)*, A0A0R1BWY9 (A. *nosocomialis)*. Residues lining the CR are coloured blue.

**Fig. S2.**
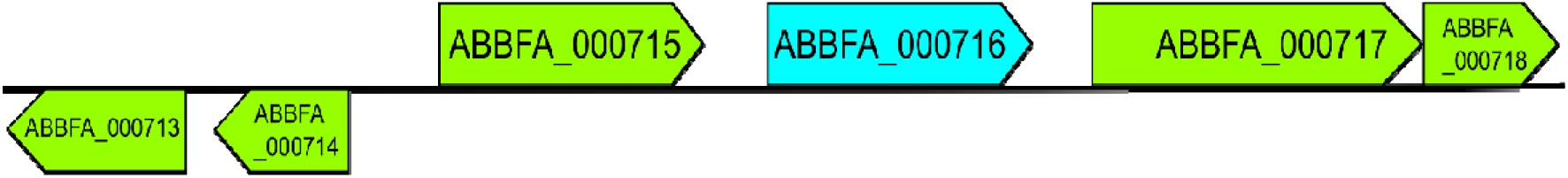
Genomic locus of ABBFA_000716 *dcaP* from *A. baumannii* from the KEGG genome database.

**Fig. S3.**
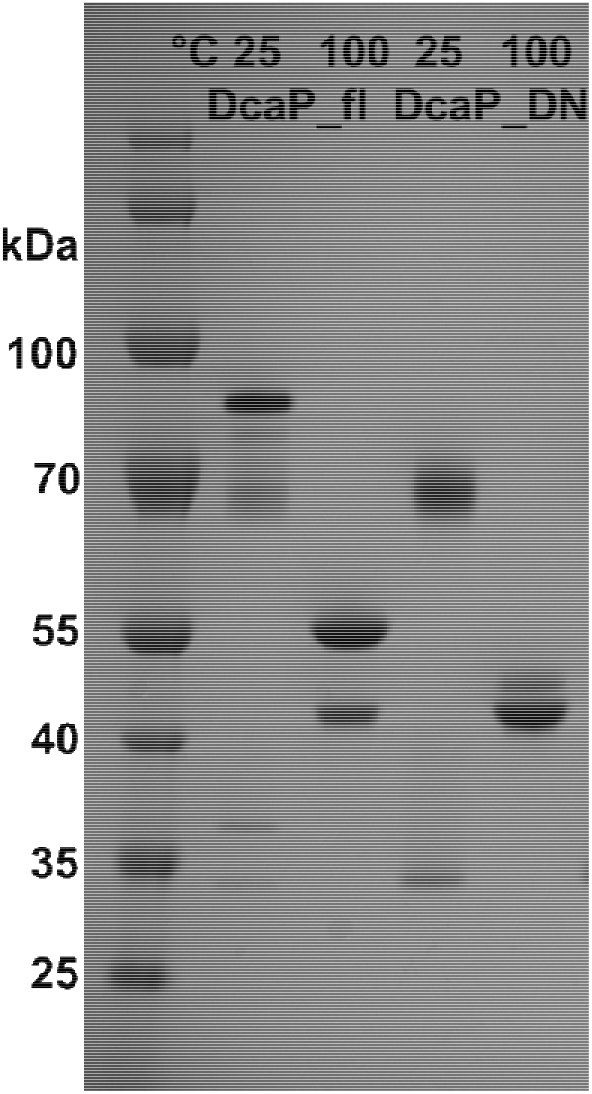
SDS-gel of DcaP constructs DcaP_fl_, and DcaP_DN40_ after the second gel filtration. Each sample is loaded as non-boiled and boiled protein sample.

**Fig. S4.**
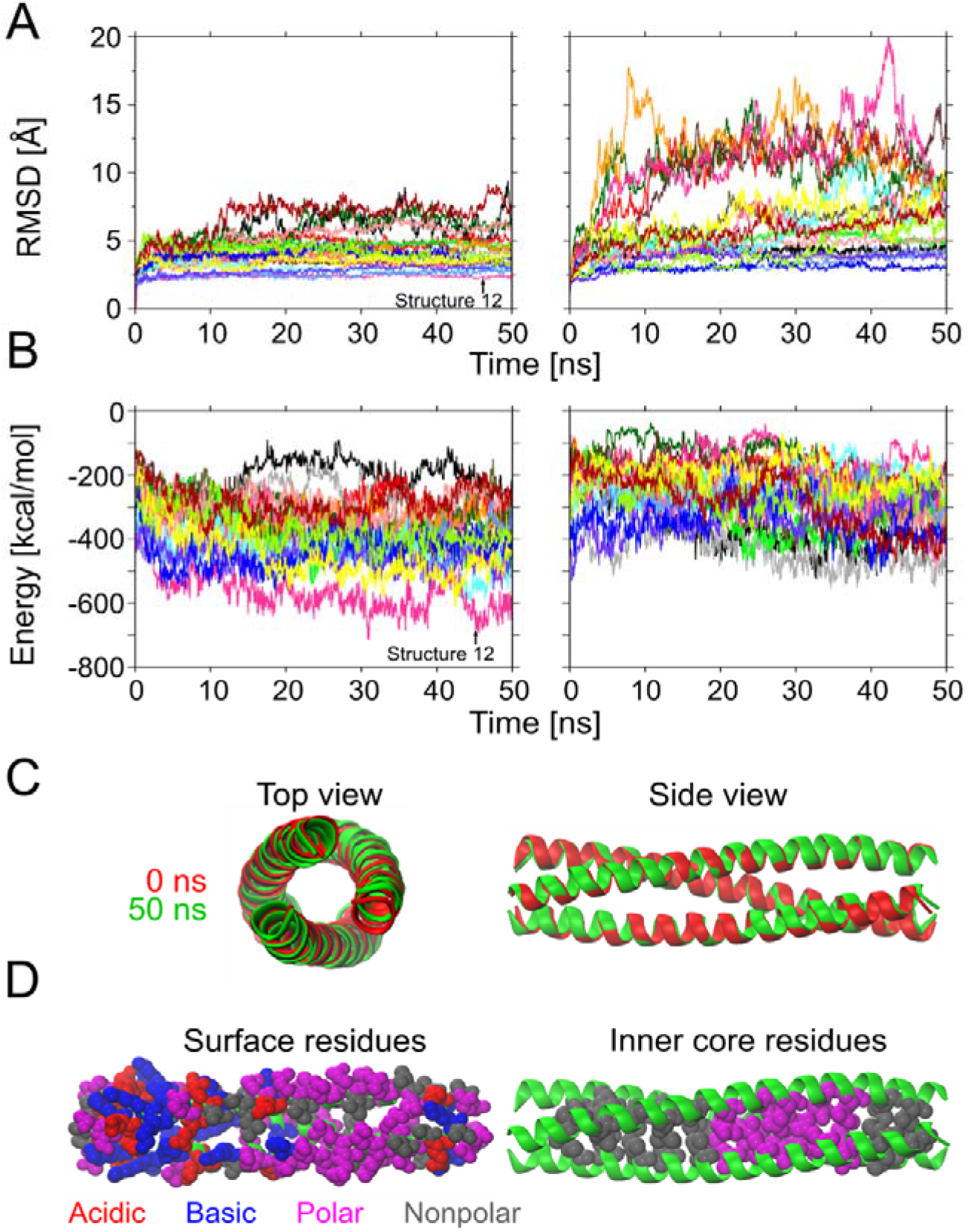
A, RMSD for all atoms of the 36 predicted coiled-coil structures (16 each in left and right panels) in the unbiased MD simulations. B, Interaction energy of one monomer with the other two monomers for all 36 predicted structures. The interaction energy includes the contributions of the electrostatic and van der Waals energy terms. C, Top and side view of the aligned structure #12 at 0 and 50 ns. D, Surface and inner core residues for structure #12. The favorable packing of the polar and nonpolar residues inside the core of the coiled-coil structure #12 cause its higher stability.

**Fig. S5.**
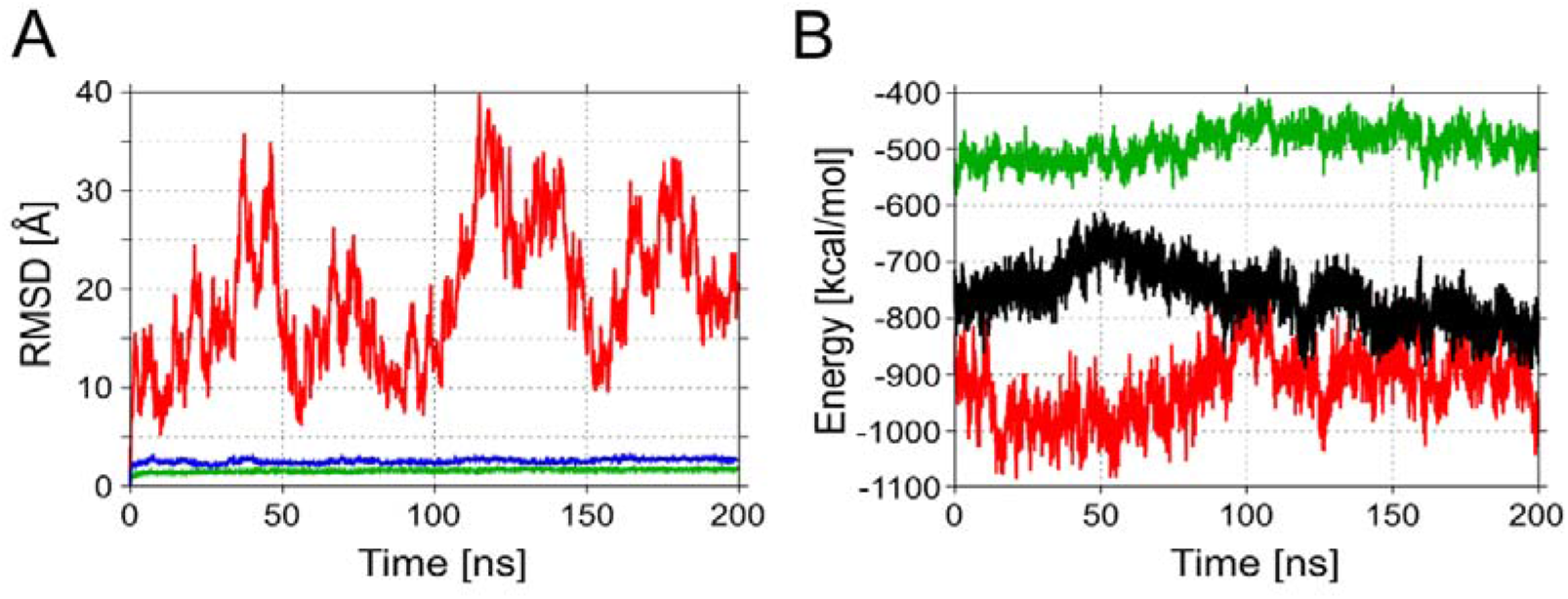
A, RMSD for all atoms of the barrel (green) and N-terminal domain (red) of DcaP_fl_ in unbiased MD simulations. The backbone atoms of the barrel residues have been aligned before the determination of the RMSD. The RMSD for all atoms of the coiled-coil structure (blue) is also shown and estimated by aligning the backbone atoms of the coiled-coil residues. B, Interaction energy of one monomer with the other two monomers for the barrel-only (green) and full length structure of DcaP (red). The interaction energy of one monomer of OmpC (black) with its two neighbouring monomers is shown for comparison.

**Fig. S6.**
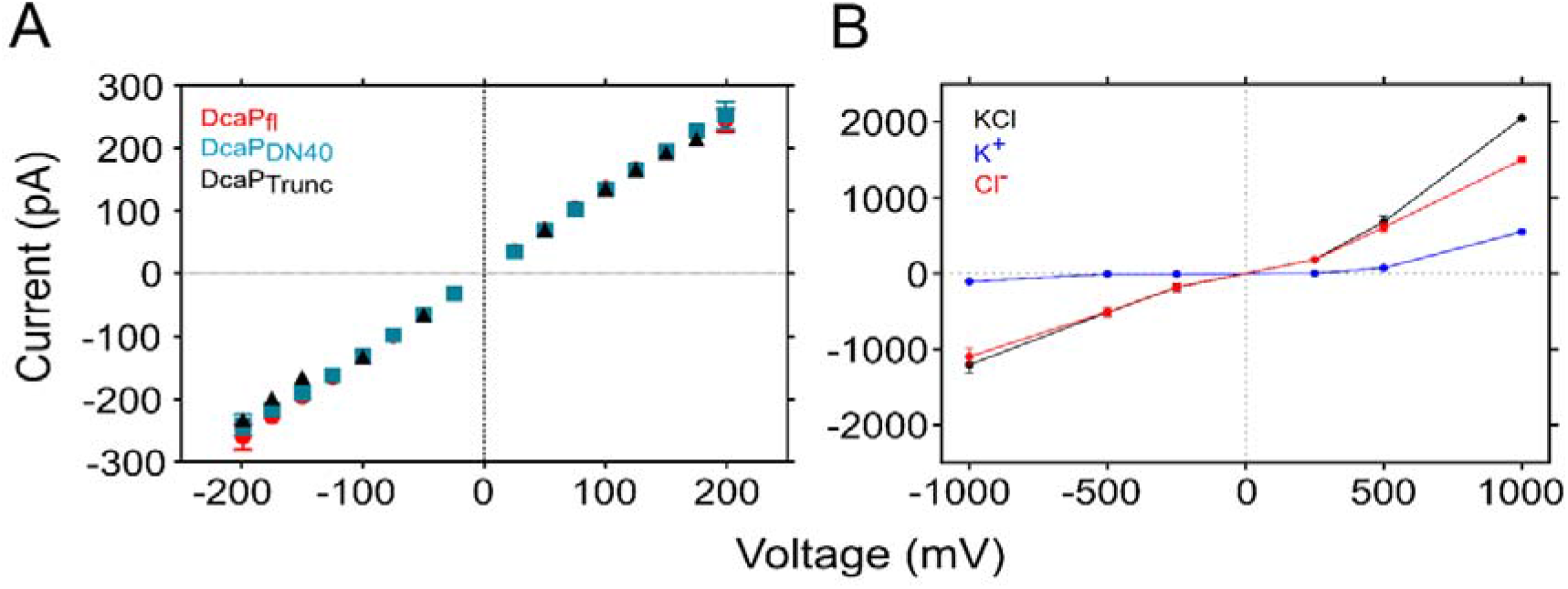
A, *I-V* curves for three the DcaP variants obtained from single channel electrophysiology experiments in 1 M KCl. B, *I-V* curve for the DcaP_Trunc_ protein calculated using applied field MD simulations in presence of 1 M KCl solution. The current generated by the individual ionic species is shown as well.

**Fig. S7.**
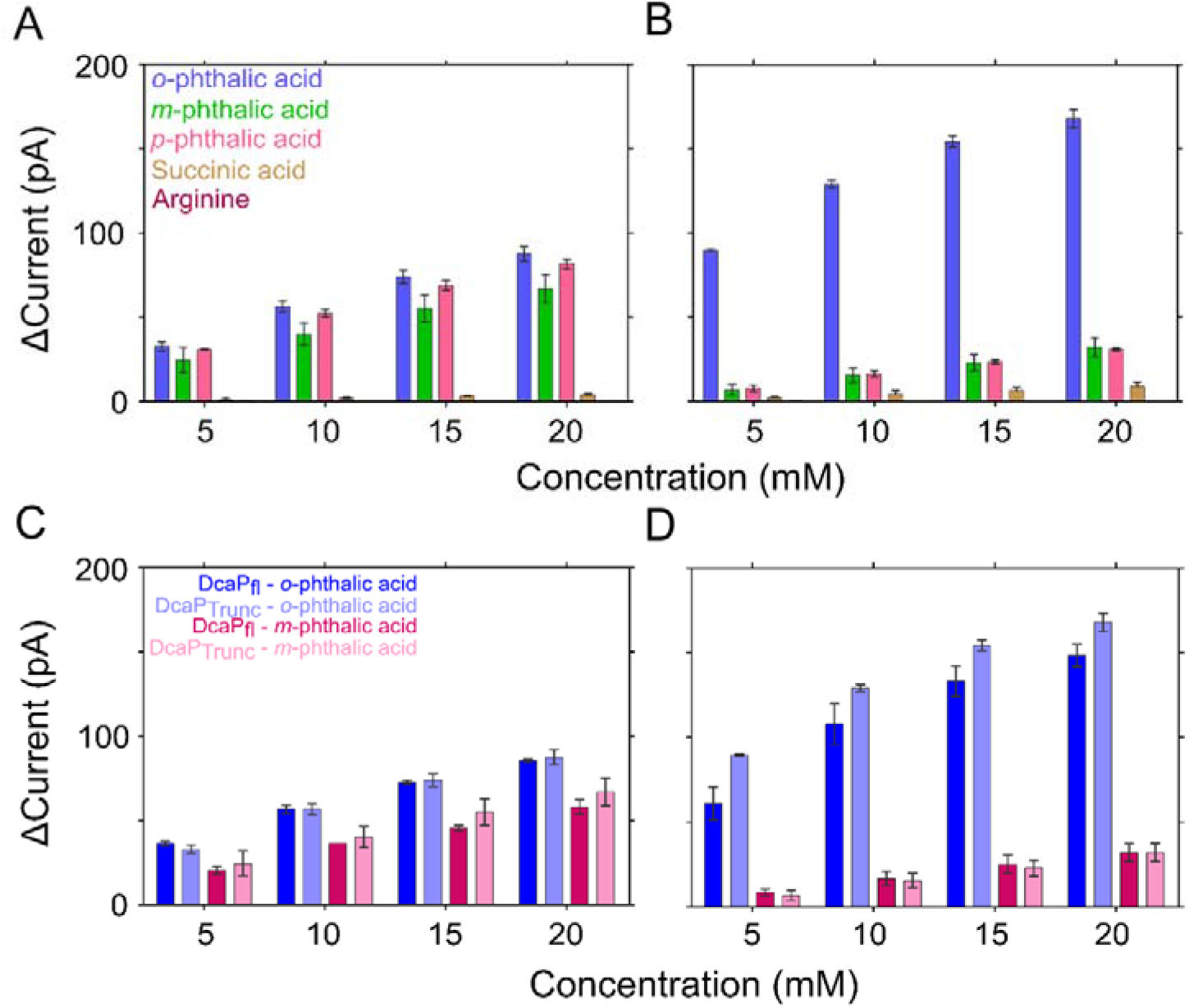
Concentration-dependent decrease in current with substrate addition for DcaP proteins. The decrease in the overall current is estimated after cis (A) and trans (B) side addition of various concentrations of *o-/m-/p*-phthalic acid, succinic acid and arginine in at +175 mV. The magnitude of the current decrease compared for DcaP_fl_ and DcaP_Trunc_ proteins after cis (C) and trans (D) side addition of *o*-phthalic acid and *m*-phthalic acid at +175 mV. The aqueous solution was buffered at 1 M KCl 10 mM HEPES pH 7.

**Fig. S8.**
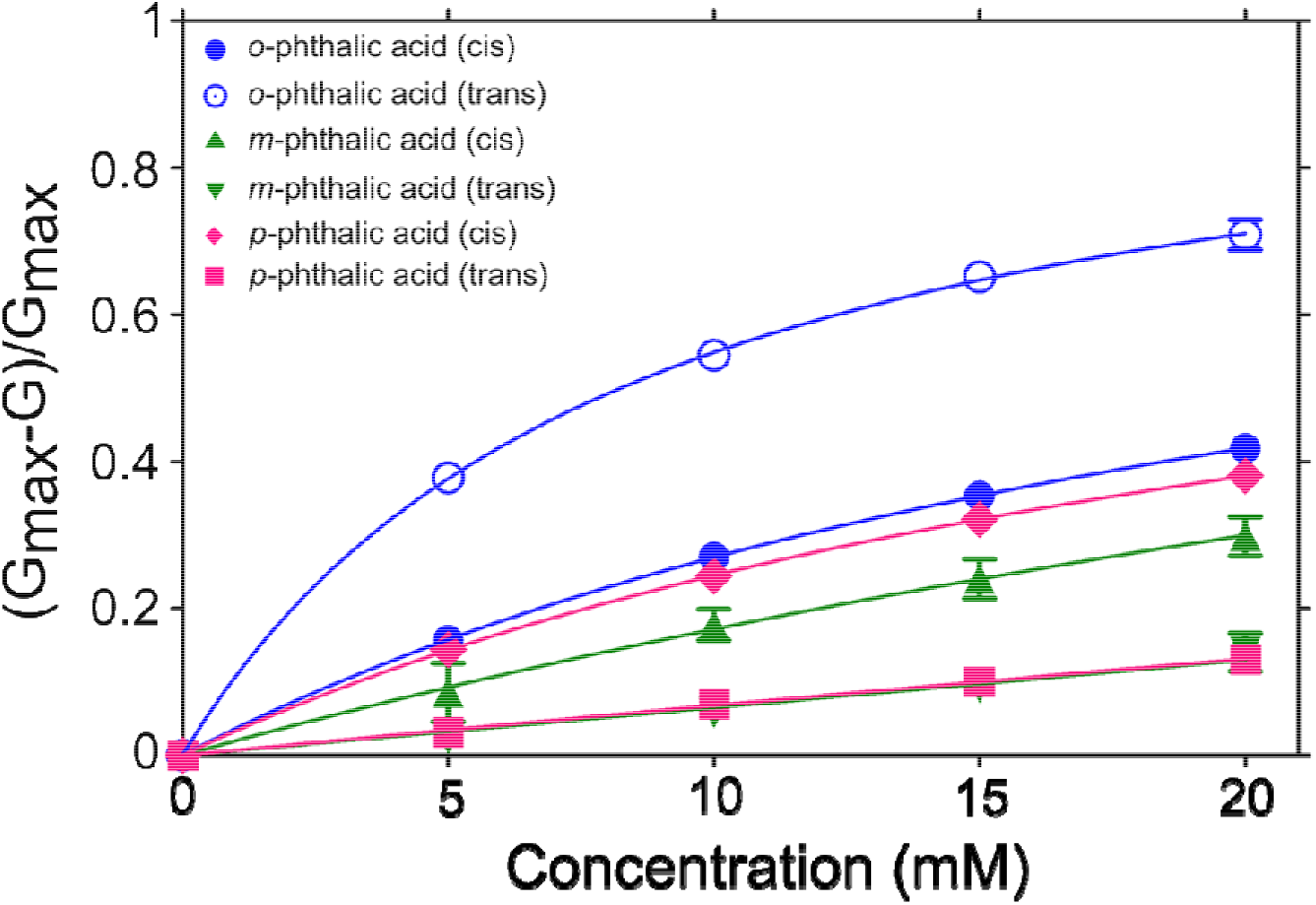
Langmuir plots of DcaP with phthalates. The binding curves of *o*-phthalic acid, *m*-phthalic acid and *p*-phtalic acid with DcaP_*Trunc*_ protein are estimated after cis or trans side addition of substrates. Binding affinities are dependent on the side of addition and type of phthalate.

**Fig. S9.**
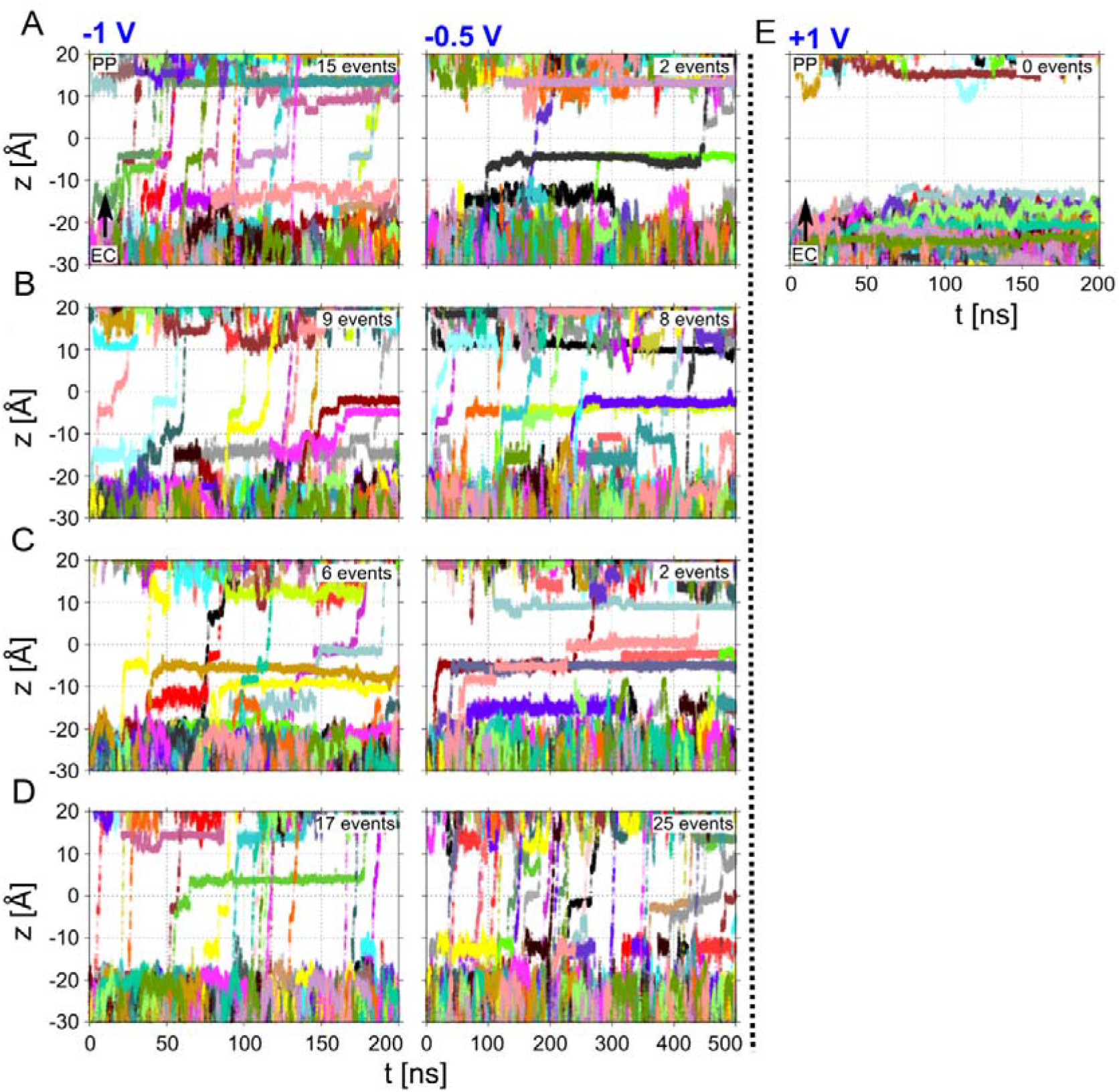
Substrate translocation in applied field MD simulations. Time evolution of the distance (2 of the substrates *o*-phthalic acid (A), *m*-phthalic acid (B), *p*-phthalic acid (C), succinic acid (D), and arginine (E), with respect to the channel during applied field MD simulations performed at −1.0 and −0.5 V. The distance *z* represents the difference along the channel axis between the center of mass of the substrate and the channel. Each substrate molecule is depicted with a different color in the individual simulations. EC and PP corresponds to extracellular and periplasmic side of the channel. The constriction region is from z = −7 to 4 Å. The number of observed translocation events are indicated for each panel.

**Fig. S10.**
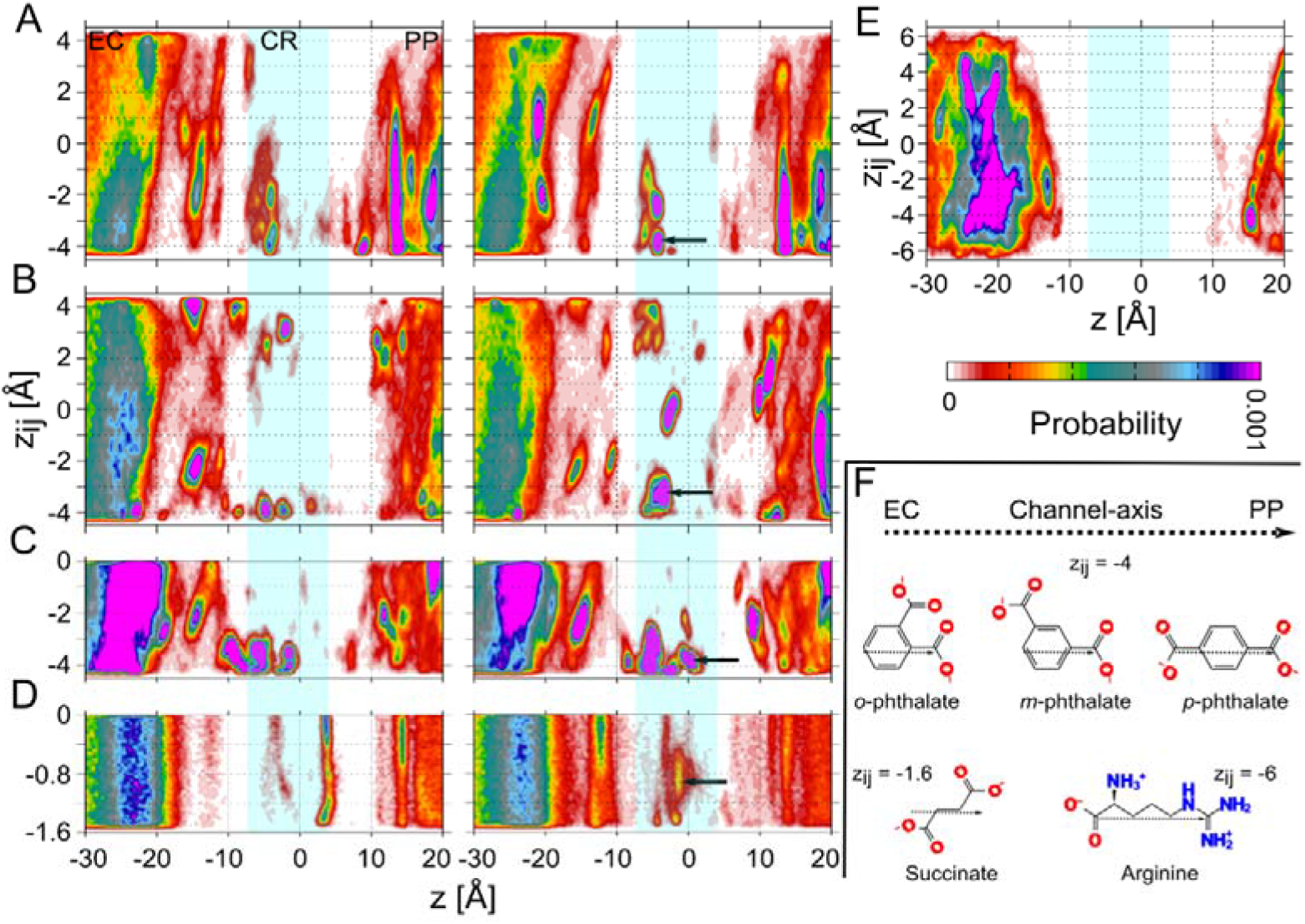
Normalized probability distribution of the substrates, *o*-phthalic acid (A), *m*- phthalic acid (B), *p*-phthalic acid (C), succinic acid (D), and arginine (E), with respect to *z* (distance along the channel axis) and *z_ij_* (orientation with respect to channel axis) during the applied field MD simulations. F, Each molecule is shown in the orientation with respect to the channel axis for the minimum values of *z_ij_*. The orientation *z_ij_* represents the *z*-component difference of the two atoms connected by the dotted arrow on each molecule. The values of *z_ij_* from the minimum to maximum can be mapped to 180 - 0° in terms of the angular variable (θ) for *o*-phthalic acid, *m*-phthalic acid, and arginine using the dot product equation 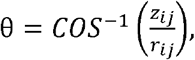 where *r_ij_* is the norm of the vector connecting the same atoms used for *z_ij_*. For example, *z_ij_* ϵ [−4,4] Å is equivalent to θ ϵ [180,0]° for *o*-phthalic acid. In case of *p*-phthalic acid and succinic acid, the range of the *z_ij_* values is limited from minimum to 0 due to symmetry of the molecules and can be mapped to 180 - 90°. The arrows in the first four panels indicate the most favorable orientations of the substrates in the constriction region and are shown in Fig. 4 of the main text.

**Fig. S11.**
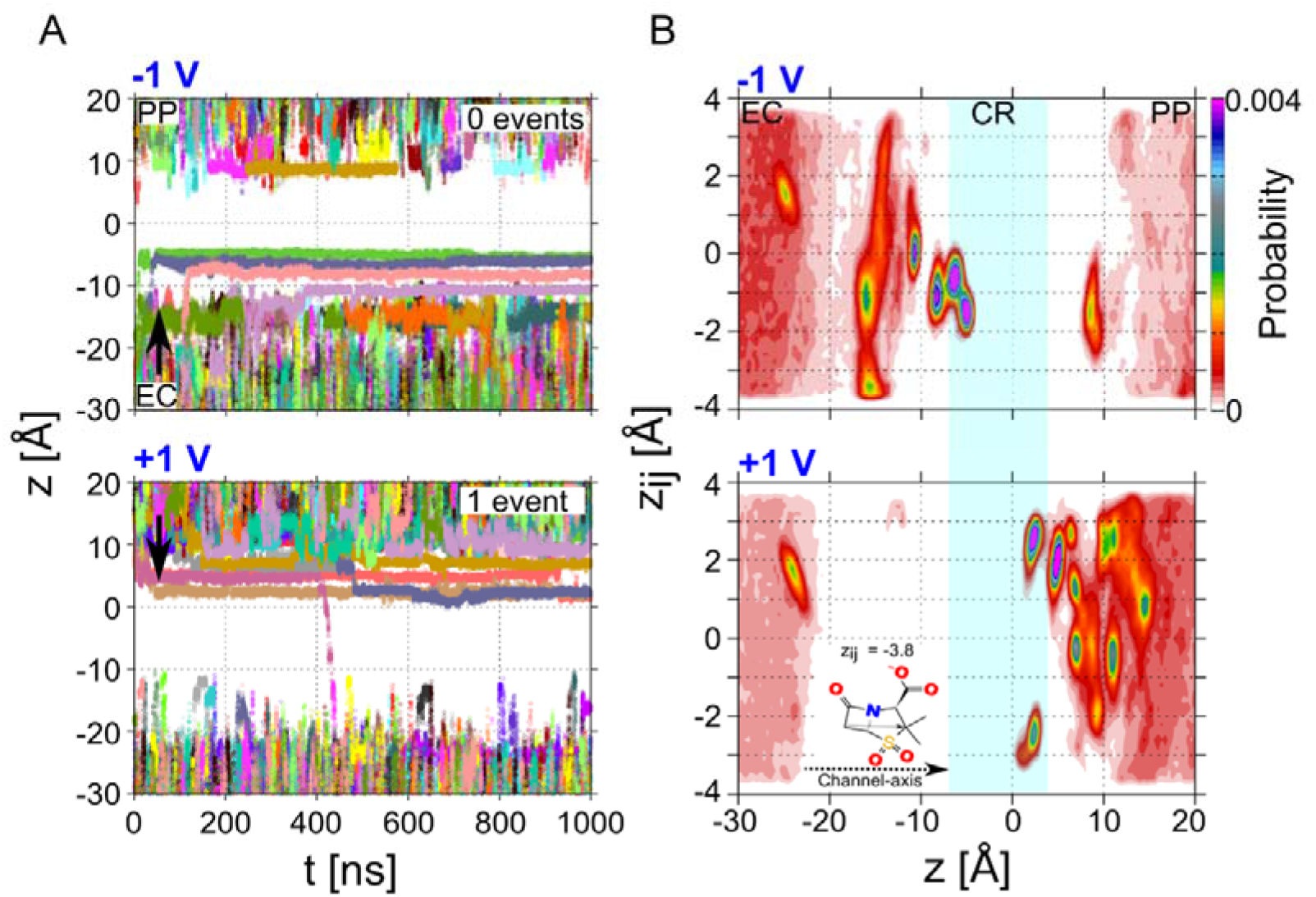
A, Time evolution of the distance (*z*) of sulbactam with respect to the channel during the applied field MD simulation performed at −1.0 (upper panel) and +1.0 V (lower panel). B, The distribution of the orientation *(z_ij_*) of the sulbactam molecule is shown with respect to the distance (*z*). Sulbactam is shown in the orientation with respect to the channel axis for the minimum values of *z_ij_* (lower panel). The two atoms used for the calculation of the z-component difference are connected by the dotted arrow. The values of *z_ij_* from the minimum to maximum can be mapped to 180 - 0° in terms of the angular variable (θ) as explained in Supplementary Fig. 10.

**Fig. S12.**
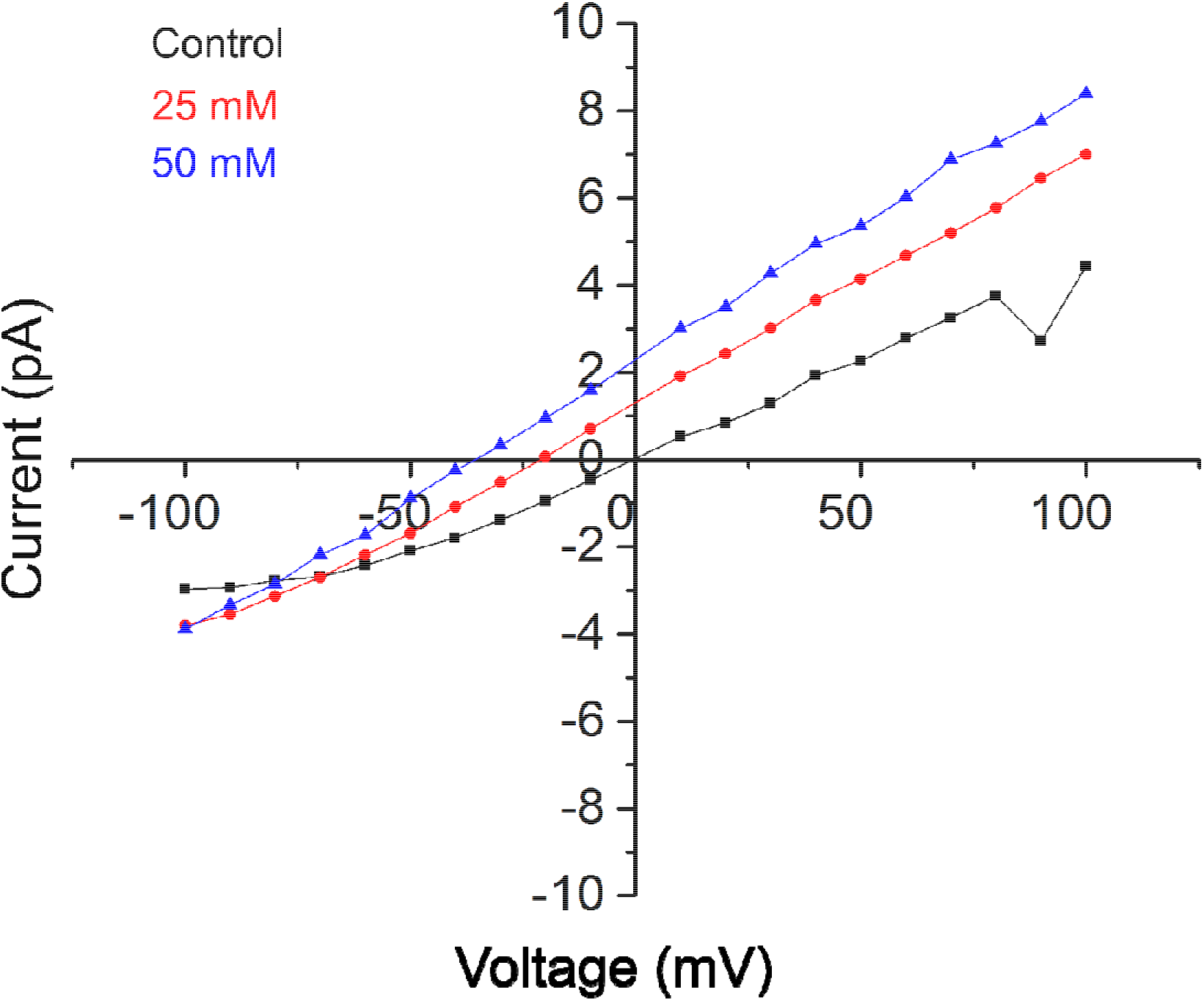
Reversal potential measurements of sulbactam, indicating a shift in the voltage on addition of the antibiotic on the cis side at 25 mM and 50 mM in 10 mM KCl 1 mM HEPES pH 7.

**Fig. S13.**
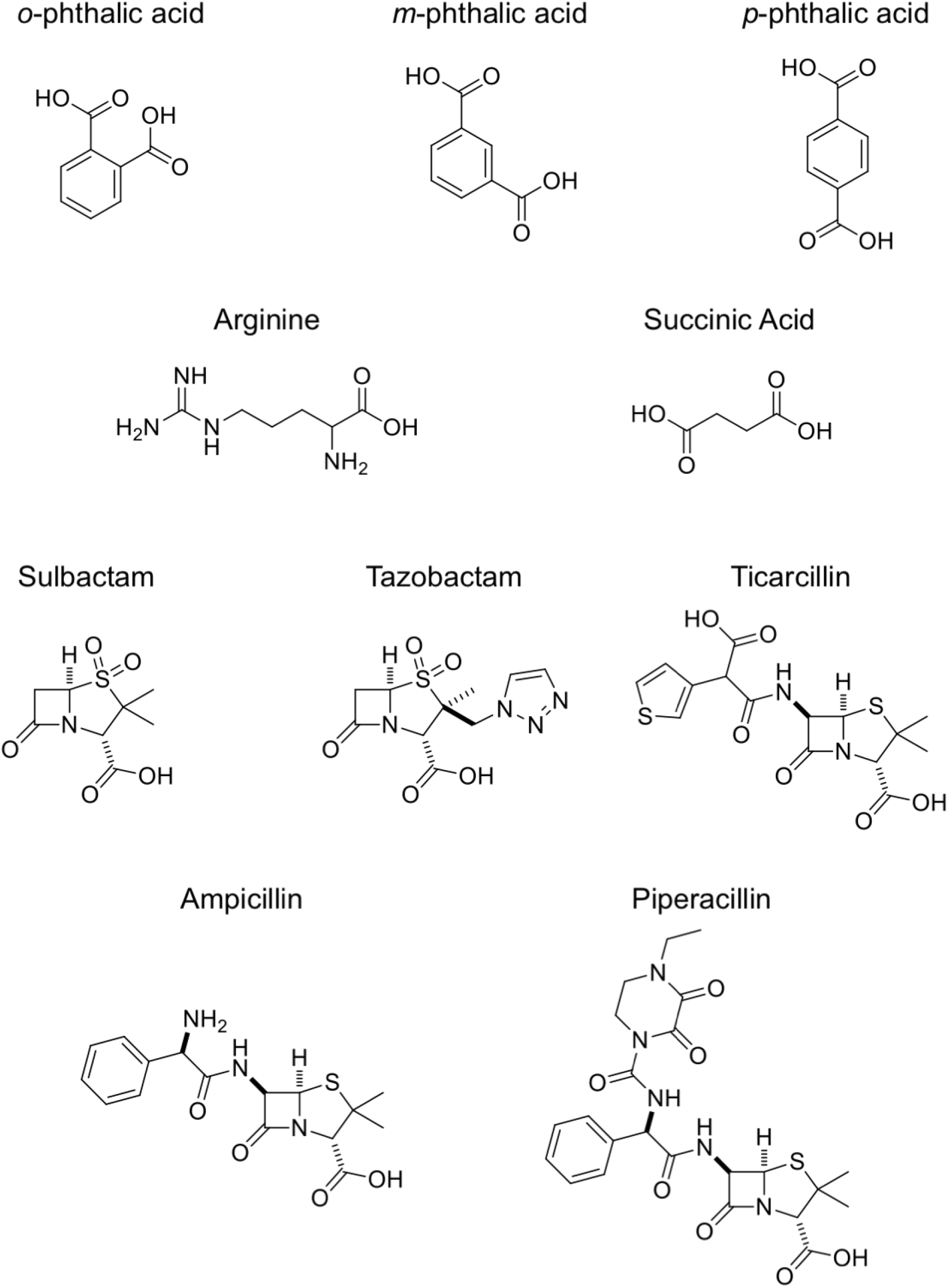
Structures of substrates and antibiotics.

**TABLE S1.**
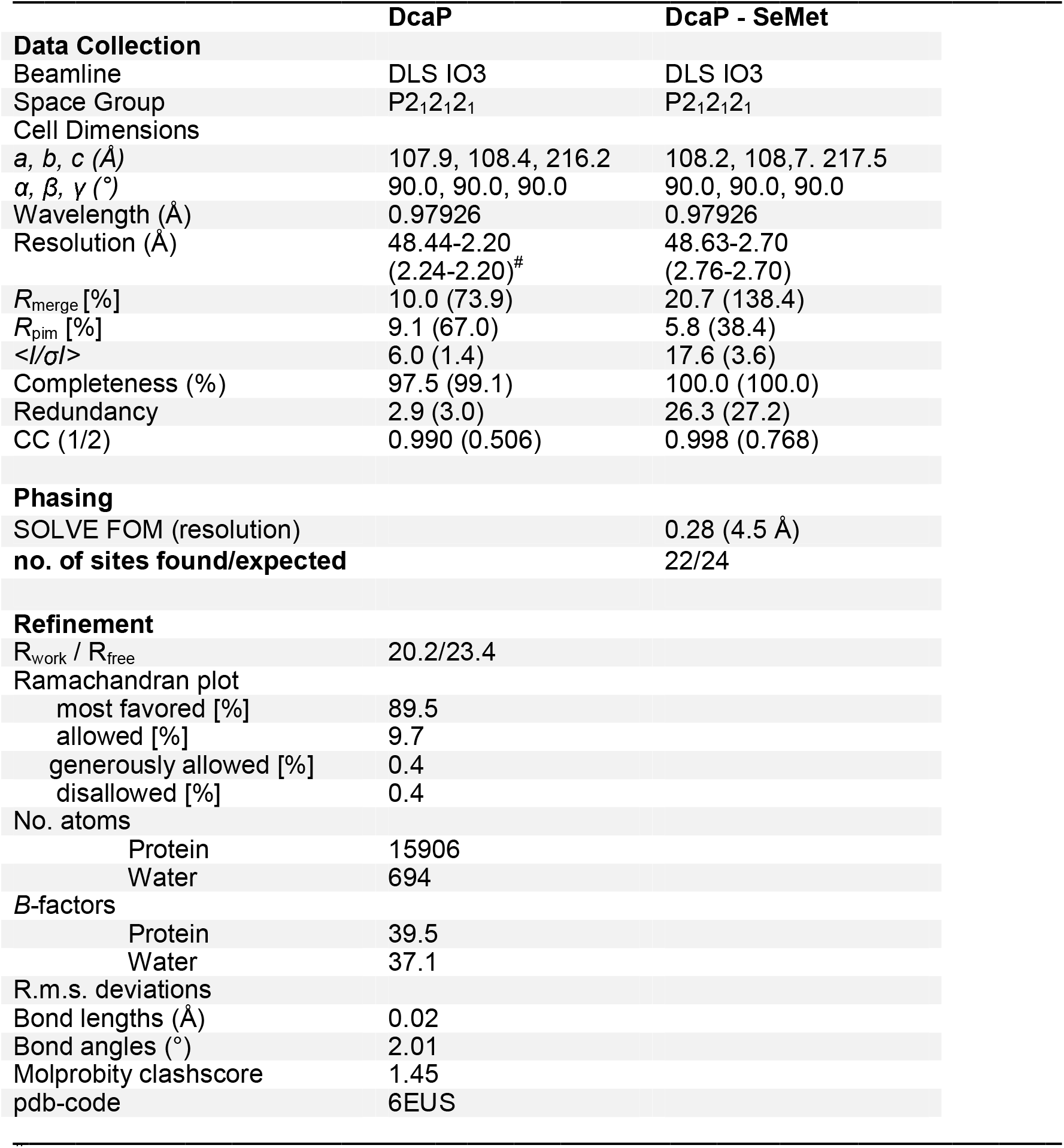
Data Collection and Refinement statistics for DcaP_DN40_.

^#^Values in parentheses are for the highest resolution shell

**TABLE S2.**
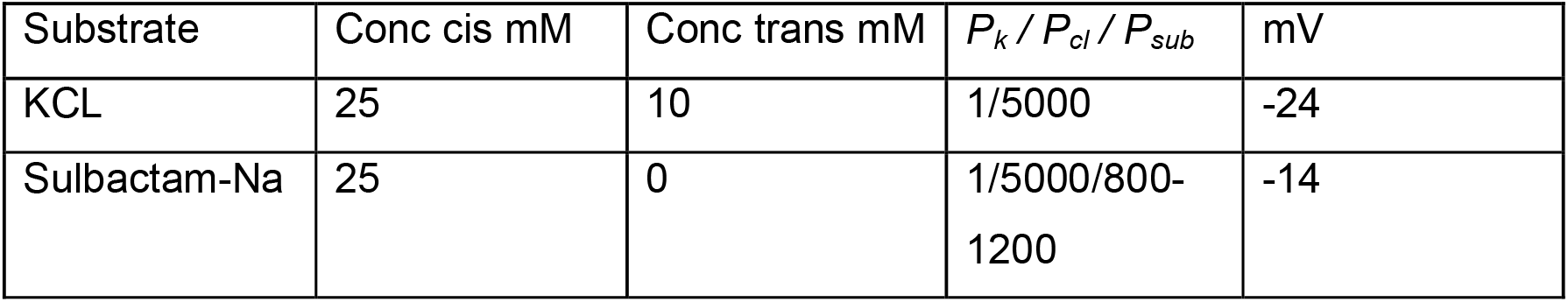
Permeability ratio and reversal potential measured after asymmetric addition of KCl or sulbactam to one side of the membrane. The concentration gradient favours the flux of negative ions and creates a negative potential.

